# Predicting modular functions and neural coding of behavior from a synaptic wiring diagram

**DOI:** 10.1101/2020.10.28.359620

**Authors:** Ashwin Vishwanathan, Alexandro D. Ramirez, Jingpeng Wu, Alex Sood, Runzhe Yang, Nico Kemnitz, Dodam Ih, Nicholas Turner, Kisuk Lee, Ignacio Tartavull, William M. Silversmith, Chris S. Jordan, Celia David, Doug Bland, Mark S. Goldman, Emre R. F. Aksay, H. Sebastian Seung, the Eyewirers

## Abstract

How much can connectomes with synaptic resolution help us understand brain function? An optimistic view is that a connectome is a major determinant of brain function and a key substrate for simulating a brain. Here we investigate the explanatory power of connectomics using a wiring diagram reconstructed from a larval zebrafish brainstem. We identify modules of strongly connected neurons that turn out to be specialized for different behavioral functions, the control of eye and body movements. We then build a neural network model using a synaptic weight matrix based on the reconstructed wiring diagram. This leads to predictions that statistically match the neural coding of eye position as observed by calcium imaging. Our work shows the promise of connectome-based brain modeling to yield experimentally testable predictions of neural activity and behavior, as well as mechanistic explanations of low-dimensional neural dynamics, a widely observed phenomenon in nervous systems.

## Introduction

It has become a truism that the connectome is “necessary but not sufficient” for understanding the brain (Bargmann, 2012; Briggman and Bock, 2012). This innocuous assertion includes a range of divergent views. Some declare that the connectome is “completely insufficient” (Nassim, 2018) and only a weak constraint at best. Others argue that the connectome is a major determinant of brain function (Denk et al., 2012), and the proper starting point for brain simulation (Seung, 2012). Due to technological advances in connectomics (Lee et al., 2019; Xu et al., 2020), the debate is moving into the domain of empirical study. Here we investigate the power of connectome-based modeling of brain function using a wiring diagram reconstructed from a larval zebrafish brainstem.

We reconstructed the synaptic connectivity of 3000 neurons from a 3D electron microscopic image. Despite highly intermingled cell bodies and arbors, the high resolution of the data enables graph clustering algorithms to identify two modules with stronger connectivity within than between modules. The modules are biologically validated using information about known cell types, and turn out to be specialized for different behaviors: one for eye movements and the other for body movements. The oculomotor module is in turn subdivided into two submodules, which appear to be specialized for movements of the two eyes. Our linking of structure and behavior by modularity analysis at synaptic resolution is novel in the vertebrate nervous system. It is especially remarkable in the brainstem reticular formation, which was regarded by classical anatomists as “undifferentiated” (Allen, 1932) or “diffuse” (Ramón-Moliner and Nauta, 1966). Previous findings of functionally specialized modules by connectivity analysis at synaptic resolution were confined to invertebrate nervous systems (Varshney et al., 2011; Jarrell et al., 2012; Pavlovic et al., 2014).

Having found modules, we then consider whether the wiring diagram can elucidate more detailed aspects of oculomotor function. When relating the wiring diagram to function, a common approach starts by extracting rules of synaptic connectivity between neuronal cell types, and then applies these rules to explain function (Seung, 2009). This approach has worked well for direction selectivity in the retina (Briggman et al., 2011; Kim et al., 2014; Ding et al., 2016), but is not guaranteed to generalize to other vertebrate brain structures that seem less stereotyped and precisely organized than the retina. Here we bypass connectivity rules and take a more direct approach: use the wiring diagram to estimate a synaptic weight matrix characterizing physiological interactions between neurons, and literally insert that matrix into a network model incorporating a minimal number of additional constraints from physiology.

Although the approach is naive, we use it to create a network model that predicts how eye position is encoded in neural activity. There are numerous reasons why our approach might have failed. We estimate the physiological strength of a connection based on the number of synapses involved, which is a rather crude measure. The model largely neglects the complexities of nonlinear dendritic integration, neuromodulation, and many other aspects of biophysics and biochemistry of neurons (Bargmann and Marder, 2013). Surprisingly, the predictions turn out to be statistically consistent at a population level with neural activity recorded by calcium imaging of larval zebrafish during oculomotor behavior. As far as we know, this is the first time that a vertebrate wiring diagram has been used to create a neural network model that predicts the encoding of behavior in neural activity. This success shows the promise of connectome-based brain modeling, an approach that should become even more powerful as more biophysical realism is incorporated.

The oculomotor system is of broad conceptual interest because it is a classic example of low dimensional neural activity dynamics (Seung, 1996). Low dimensional dynamical attractors have been proposed to underlie a large range of computations, from storage of working memory to reduction of noise in sensory and cognitive representations (Yoon et al., 2013; Daie et al., 2015; Kim et al., 2017; Green et al., 2017). However, it remains mysterious how these attractors are created at the level of individual cells and circuits. Furthermore, little attention has been paid to what features of the circuitry lead to the observed low-dimensional patterns of activity in recurrently connected networks and their downstream targets. Our reconstructed wiring diagram allows us to address these questions and thereby provides unprecedented insights into the cellular resolution circuitry governing the generation and transmission to target structures of low-dimensional attractors hypothesized to underlie a broad range of cognitive and non-cognitive tasks.

## Results

### Neuronal wiring diagram reconstructed from a vertebrate brainstem

We applied serial section electron microscopy (EM) to reconstruct synaptic connections between neurons in a larval zebrafish brainstem (Figure 1). The imaged dataset targeted a volume that is known to include neurons involved in eye movements (Schoonheim et al., 2010; Miri et al., 2011a; Daie et al., 2015; Lee et al., 2015; Vishwanathan et al., 2017). First, the dataset of Ref. (Vishwanathan et al., 2017) was extended by additional imaging of the same serial sections. By improving the alignment of EM images relative to our original work, we created a 3D image stack that was amenable to semiautomated reconstruction (Lee et al., 2019) of 100× more neurons than before. We trained a convolutional neural network to detect neuronal boundaries (Methods) (Lee et al., 2019), and used the output to generate an automated segmentation of the volume. To correct the errors in the segmentation, we repurposed Eyewire, which was originally developed for proofreading mouse retinal neurons (Kim et al., 2014). Eyewirers proofread ~3000 objects, which included neurons with cell bodies in the volume as well as “orphan” neurites with no cell body in the volume (Figure 1A). We will refer to all such objects as “nodes” of the reconstructed network. Convolutional networks were used to automatically detect synaptic clefts, and assign presynaptic and postsynaptic partner nodes to each cleft (Turner et al., 2020) (Figure 1B-D, Methods). The reconstructed dataset contained 2824 nodes, 44949 connections between pairs of nodes, and 75163 synapses. Most connections (65%) involved just one synapse, but some involved two (19%), three (7.9%), four (3.7%), or more (4.0%) synapses, up to a maximum of 21 (Figure S1A).

**Figure 1:**
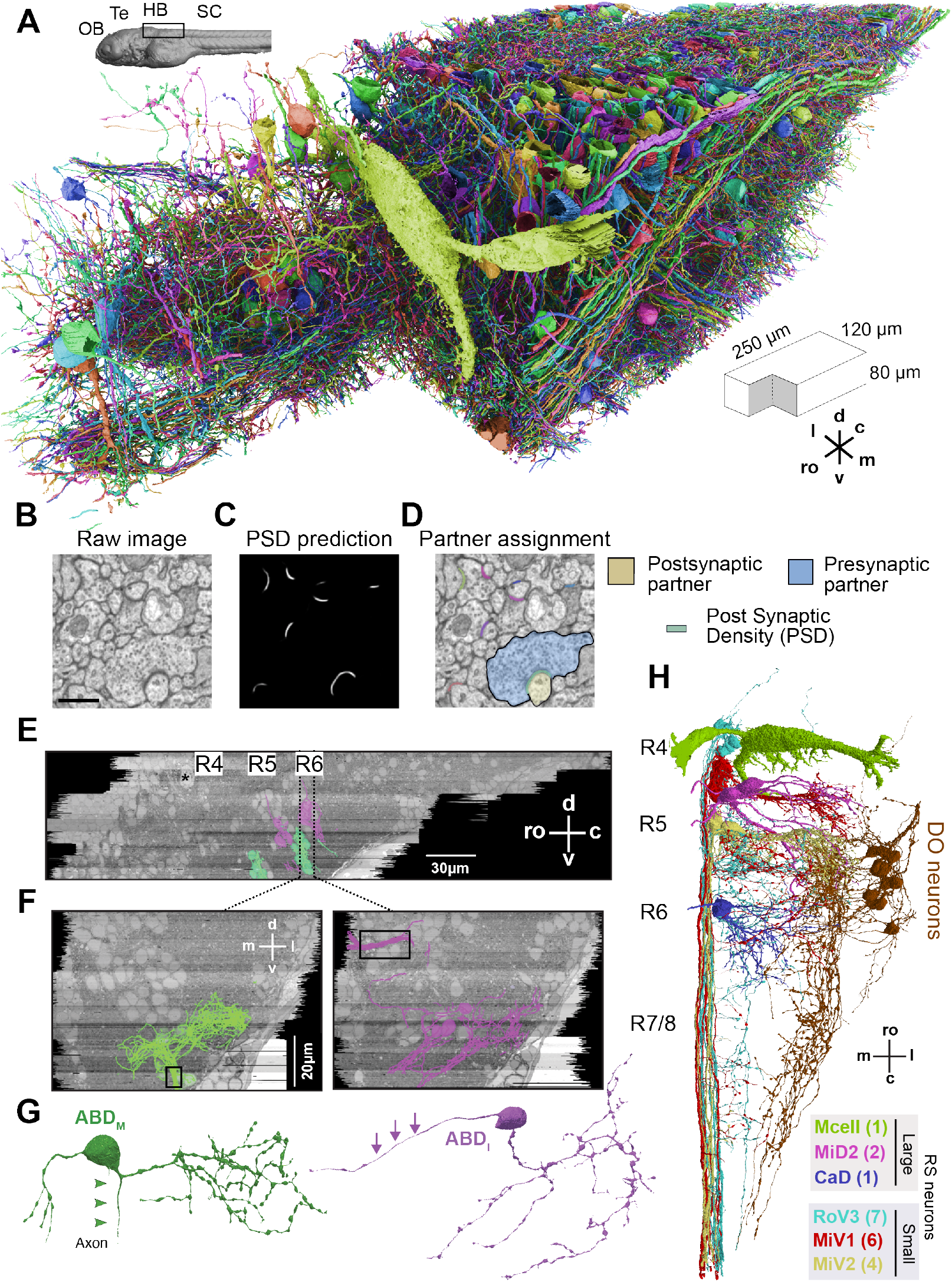
EM reconstructions of brainstem neurons. **A.** 3D rendering of reconstructed neurons. Large green cell body in the foreground is the Mauthner neuron (ro - rostral; c - caudal; d - dorsal; v - ventral; l - lateral; m - medial). Inset (top left) shows location of the unilateral EM volume (black box) relative to the olfactory bulb (OB), tectum (TE), hindbrain (HB), and spinal cord (SC). **B-D.** Automatic synapse detection and partner assignment. B) Raw EM image. Scale bar is 750nm. C) Postsynaptic densities (PSDs) identified by a convolutional net. D) PSDs (red) overlaid onto the original raw image, together with an exemplar presynaptic (blue) and postsynaptic (yellow) partnership identified by a second convolutional net. **E.** Sagittal view of identified abducens motor (ABD_M_, green) and abducens internuclear (ABD_I_, magenta) neurons overlaid over representative EM planes (R - rhombomere; * - Mauthner cell soma). **F.** Coronal planes showing the locations of the ABD_M_ (left) and ABD_I_ (right) neurons at the planes indicated by dotted black lines in e sagittal view. Black boxes highlight nerve bundles from these populations. **G.** Representative ABD_M_ and ABD_I_neurons with arrows indicating the axons. **H.** Reconstructions of large and small reticulospinal (RS) neurons and dorsal octaval (DO) neurons.

After registration of our EM volume to the Z-Brain reference atlas (Randlett et al., 2015) (Figure S2), several important groups of neurons and neurites were identified (Methods). It was straightforward to recognize two populations of reticulospinal (RS) projection neurons involved in escape behaviors and controlling movements of the body axis. The neurons of one population were large and dorsally located, and of the other were smaller and ventromedially located (Figure 1H) Comparison with transgenic lines and cranial nerves in the atlas yielded the abducens (ABD) neurons controlling extraocular muscles, and Descending Octaval (DO) neurons that mediate optokinetic and vestibular signals (Pastor et al., 2019) as part of the sensorimotor transformations needed to control eye movement (Figure 1E-H; Figure S3, S4; Methods).

### Axial and oculomotor modules in the brainstem

Before looking for modularity, we divided the reconstructed neurons into “center” and “periphery” (Methods). Neurons in the “center” (540 neurons) are recurrently connected to other reconstructed neurons that are expected to predominate in establishing collective dynamics. Neurons in the “periphery” (2344 nodes), in contrast, are involved in feedforward pathways that supply input to the center, transmit output from the center, or have negligible recurrent connectivity as quantified by eigenvector centrality (62 nodes). For example, the periphery includes ABD neurons, which mediate a downstream pathway to the extraocular muscles, as mentioned above. The periphery also includes most reticulospinal projecting neurons, such as the Mauthner cell, MiV1 and MiV2 cells (Figure 1H).

Complex systems, both biological (Wagner et al., 2007) and artificial (Baldwin et al., 2000), can often be divided into modules, such that interactions within modules are stronger than interactions between them. While such modules are structurally defined, they often turn out to be functionally specialized. To identify modules in our reconstructed wiring diagram, we applied a graph clustering algorithm to the center of the network (Methods). This analysis revealed a block structure in the connection matrix (Figure 2A). The diagonal blocks of the matrix contain the connections within modules, and the off-diagonal blocks contain connections between modules. The diagonal blocks are more densely connected than the off-diagonal blocks, meaning that within-module connectivity is stronger than between-module connectivity, which is what the graph clustering algorithm is meant to achieve. We designated the two modules as modA and modO, for reasons explained below.

**Figure 2:**
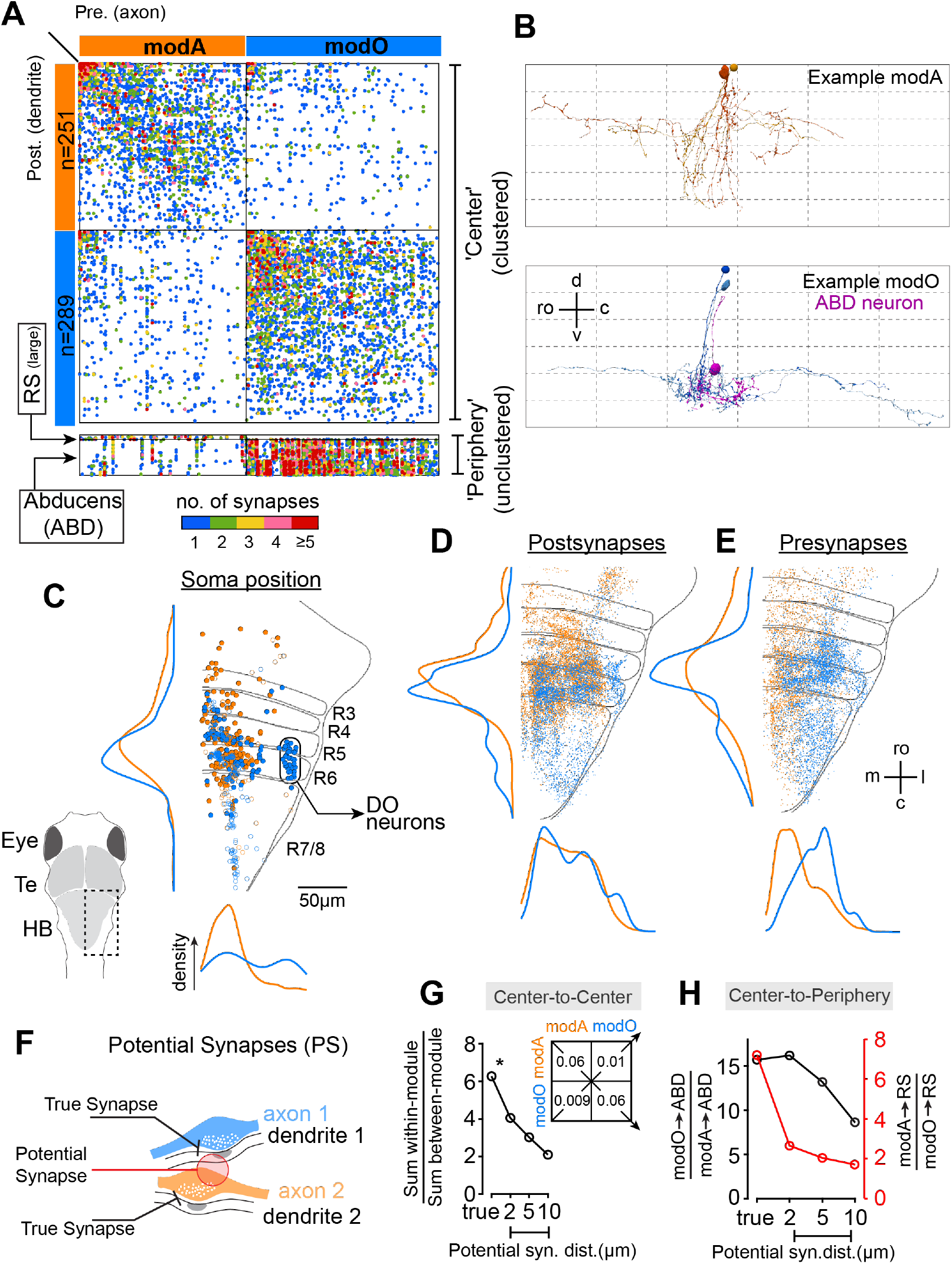
Modularity and functional specialization of interneurons. **A.** (Top) Matrix of connections in the ‘center’ of the wiring diagram, with neurons clustered into two modules (modA, modO). (Bottom) Matrix of connections from center to select reticulospinal (RS) and abducens (ABD) neurons in the periphery. **B.** (Top) Example connected pairs of modA neurons. (Bottom) Example connected pair of modO neurons (light and dark blue) and the overlap of their axons with the dendrite of an abducens internuclear cell (magenta) (ro - rostral; c - causal; d - dorsal; v - ventral). Grid in the background is the same in both panels to facilitate comparison. **C.** Locations of reconstructed neuron soma (modA - orange, modO - blue) projected onto the horizontal plane and 1D densities along the mediolateral (bottom) and rostrocaudal (right) axes (DO - descending octaval; R - rhombomere). Closed circles are neurons with complete somas inside the reconstructed EM volume. Open circles are locations of the primary neurites exiting the top of the EM volume for cells with somas above the volume. Inset cartoon shows the region of the hindbrain in the figure. Te - tectum, HB - hindbrain. **D.** Postsynapses of neurons in modA and modO along with 1D densities. Every 5^th^ postsynaptic density is plotted for clarity. **E.** Presynapses of neurons in modA and modO along with 1D densities. Every 10^th^ presynaptic terminal is plotted for clarity. (m - medial; l - lateral). **F.** Schematic illustrating the definition of a potential (i.e. false) synaptic connection identified when a presynaptic terminal (e.g. axon 2) is proximal (red) to a postsynaptic density (e.g. dendrite 1) but not actually in contact with it. **G.** Ratio of the number of within-module to the number of between-module synapses versus threshold distance for true and potential synapses. Table lists the actual true synapse (TS) numbers for data point with asterisk *. H. Ratio of numbers of synapses from neurons in modA and modO to peripheral neurons (ABD, RS).

For biological validation of the modules, we checked them against information that was not used by the clustering algorithm. Namely, we checked the relation of the modules to two neuron classes (RS and ABD) identified above (Figure 2A). RS neurons in the periphery received much stronger connectivity from modA than from modO (Figure 2A, RS; Figure S8). Of the 10 smaller RS neurons contained in the center, all were in modA, and none were in modO (Figure S8A, black arrows). Furthermore, the 15 RS neurons (small and large) contained in the periphery received much stronger connectivity from modA than from modO (Figure S8). Since the RS neurons are known to be involved in turning or swimming movements (Gahtan et al., 2002; Orger et al., 2008; Huang et al., 2013; Bhattacharyya et al., 2017), we propose that modA plays a role in movements of the body axis, and refer to it as the “axial module.”

All 54 ABD neurons were in the periphery, and received much stronger connectivity from modO than from modA (Figure 2A). All 34 of the DO neurons (Figure 2c) were members of modO; none were in modA. ABD neurons drive extraocular muscles either directly or through a disynaptic pathway. DO neurons are secondary vestibular neurons, which provide input to the vestibulo-ocular reflex. We therefore propose that modO plays a role in eye movements, and will refer to it as the “oculomotor module.”

The modules in Figure 2 were obtained with the Louvain algorithm for graph clustering (Blondel et al., 2008; Reichardt and Bornholdt, 2006; Rubinov and Sporns, 2010). Similar modules are obtained when spectral clustering (Chung, 2005) or a degree-corrected stochastic block model (Peixoto, 2014) are used (Figure S7). As we will see later, the binary division into two modules is the first step in a hierarchical clustering procedure. (For comparison with flat clusterings, see Figure S5.)

It does not appear that modA and modO can be regarded as traditional brain “nuclei,” because their somas are often intermingled and are highly distributed, extending rostrocaudally over all rhombomeres in the EM volume (Figure 2C). Postsynapses and presynapses are even more diffusely distributed than somas (Figure 2D, E). However, presynapses of modA and modO do exhibit some spatial segregation along the mediolateral axis (Figure 2E), which reflects an underlying spatial organization of axonal arbors (data not shown). Therefore we decided to probe to what extent spatial organization could be contributing to modularity.

To quantify modularity, we defined an index of wiring specificity as the ratio of the sum of within-module synapse densities to the sum of between-module synapse densities. Here synapse density is defined as the number of synapses normalized by the product of presynaptic and postsynaptic neuron numbers. The wiring specificity index was roughly 6 for the center neurons (Figure 2G), based on actual synapses. We defined a “potential synapse” as a presynapse and a postsynapse within some threshold distance of each other (Figure 2F), similar to the conventional definition as an axo-dendritic apposition within some threshold distance (Stepanyants and Chklovskii, 2005). Then we computed the wiring specificity index based on potential synapses rather than actual synapses. The index dropped to less than 3 for potential synapses defined by a distance threshold of 5μm, and close to 2 at a distance threshold of 10μm (Figure 2G, center-to-center).

The implication is that the division into modA and modO can be explained by spatial organization, as long as location information is precise to within a few microns or less (Motta et al., 2019). On the other hand, the coarse (tens of microns) mediolateral segregation of presynapses evident in Figure 2E must contribute little to modularity.

We similarly defined an index of wiring specificity for peripheral populations as the synapse density from preferred partner in the center divided by the synapse density from non-preferred partner in the center. This wiring specificity index decreases greatly for peripheral RS neurons when potential synapses are considered, but the decrease is more modest for ABD neurons (Figure 2H, center-to-periphery). We also found some statistical differences between modA and modO in total arbor length, synapse size, and synapse distance from soma (Figure S6A, B).

To further validate our claims regarding function, we performed calcium imaging throughout the hindbrain in a separate set of 20 age-matched animals. We resorted to comparison with other animals because our calcium imaging of the neurons in the EM volume (Vishwanathan et al., 2017) had highly incomplete coverage due to technical limitations. Recordings were obtained while the animals performed spontaneous eye movements in the dark, with activity ranging from neurons that exhibited bursts during saccades to those with perfectly persistent firing during fixations (Ramirez and Aksay, 2018). The images from 20 age-matched animals were combined by registering them to a reference atlas (Randlett et al., 2015). We extracted a map for eye movement signals (Methods) that was complete, in the sense that each hindbrain voxel was covered by at least three fish (Methods). Relative to modA somas, modO somas were more than twice as likely to be neighbors with somas in the map of eye movement signals (Figure S9). While this preference is not extremely strong, it seems reasonably strong given that there is intermingling of modA and modO somas (Figure 2C), and registration to the reference atlas is unlikely to remove all soma location variability across individuals. The preference disappeared when soma locations were artificially jittered by more than 10μm (Figure S9).

### Submodules of the Oculomotor network

There is a rich repertoire of oculomotor behaviors, which vary in speed of movement, patterns of binocular coordination, and other properties. Motivated by this functional diversity, we applied the same graph clustering algorithm employed in Figure 2A to modO. Doing this revealed the presence of two submodules characterized by strong within-submodule connectivity and weak between-submodule connectivity (Figure 3A).

**Figure 3:**
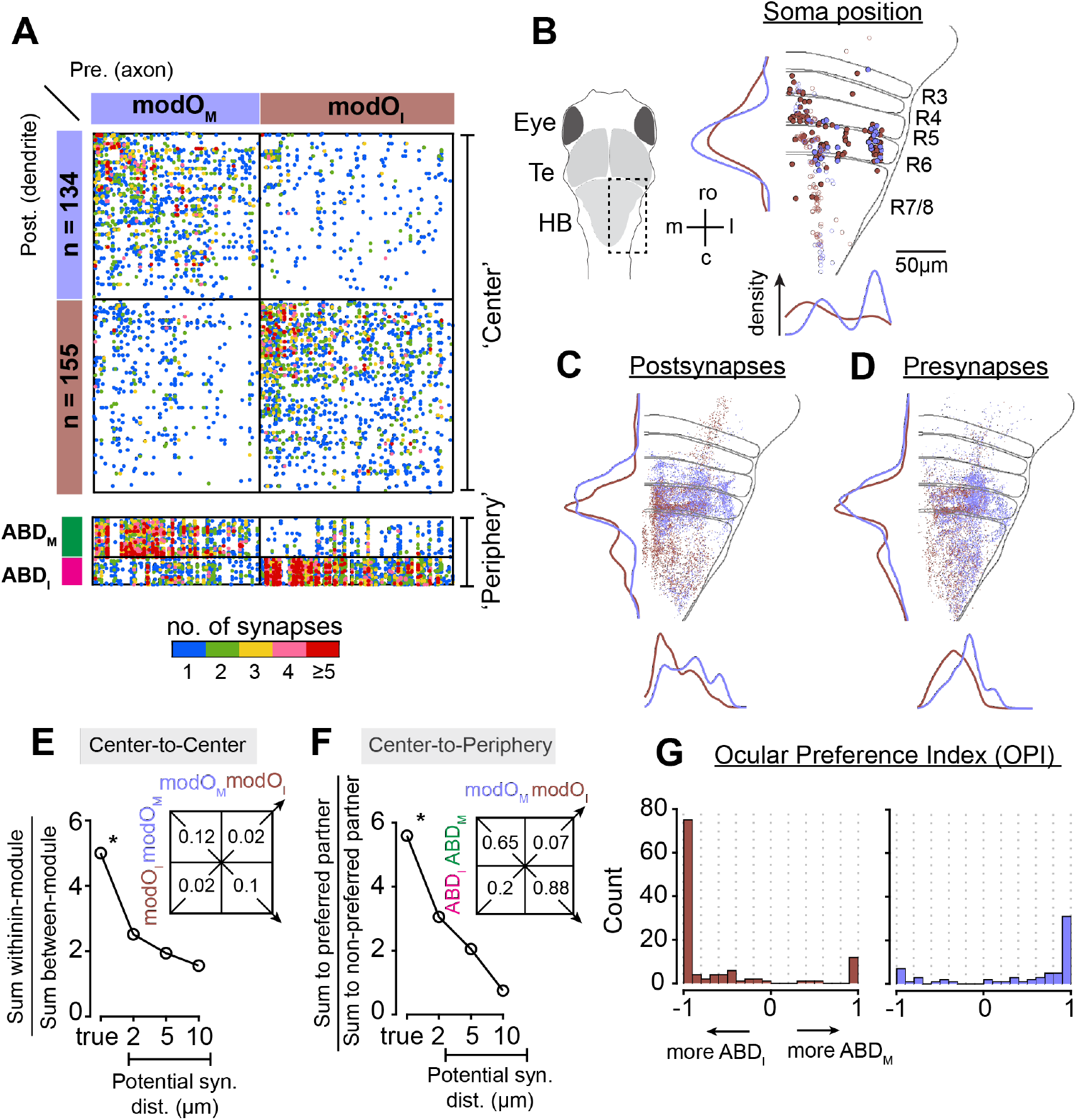
Submodules specialized for the two eyes. **A.** (Top) Matrix of connections within modO organized into two sub-modules termed modO_M_ and modO_I_ (Bottom) Projections of modO_M_ and modO_I_ onto abducens motor neurons (ABD_M_) and abducens internuclear neurons (ABD_I_) populations. **B.** Locations of reconstructed neuron soma along with 1D densities for neurons within modO_M_ (blue) modO_I_ (brown). Symbols, inset, and orientation as in Figure 3C. **C-D.** Postsynaptic densities (C) and presynaptic terminals (D) in modO_I_ and modO_M_. Every 5^th^ synaptic site was plotted for clarity. **E.** Ratio of the number of synapses within-modules modO_M_ and modO_I_ to the number between-modules as a function of potential synapses distance. Table lists the actual true synapses numbers for data point with asterisk *. **F.** Ratio of the number of synaptic contacts between a modO submodule and its preferred peripheral partner vs those between a modO submodule and its non-preferred peripheral partner. Numbers in tables represent normalized synapse counts defined as the ratio of sum of all synapses in a block to the product of the number of elements in the block. **G.** Ocular Preference index (OPI) for modO_M_ and modO_I_ neurons. DO vestibular neurons were not included.

For biological validation of the submodules, we examined the connectivity from modO to ABD neurons. This center-periphery connectivity provides independent validation, because the graph clustering algorithm relied only on intra-center connectivity. The abducens complex is composed of two groups, the motor neurons (ABD_M_) that directly drive the lateral rectus muscle of the ipsilateral eye, and the abducens internuclear neurons (ABD_I_) that indirectly drive the medial rectus muscle of the contralateral eye through a disynaptic pathway. Increased activities in both ABD_M_ and ABD_I_ neurons drive eye movements toward the side of the brain on which the neurons reside (‘ipsiversive’ movements). Neurons in one submodule of modO preferentially connected to ABD_M_ neurons, while neurons in the other submodule preferentially connected to ABD_I_ neurons. This is evident from both visual inspection (Figure 3A) and quantitative analysis (Figs. 3A, F). We therefore refer to the submodules as motor-targeting (modO_M_) and internuclear-targeting (modO_I_), and suggest they are largely involved in controlling movements of the ipsilateral and contralateral eye, respectively.

The spatial layout of somas, postsynapses and presynapses for these submodules is shown in Figure 3B-D. We again quantified the contribution of spatial organization to wiring specificity using the potential synapse formalism, and found that the division of modO into modO_M_ and modO_I_ requires spatial precision of a few microns (Figure 3E, center-to-center). A similar finding holds for wiring specificity from modO to ABD (Figure 3F, center-to-periphery). Preferences of individual neurons can be extreme: many modO cells connect to ABD_M_ only or ABD_I_ only (Figure 3G). There were some statistical differences between modO_M_ and modO_I_ in synapse size and synapse distance from the soma (Figure S6D, E).

### Predicting neural coding of eye position from the wiring diagram

We hypothesized that modO contains the “neural integrator” for horizontal eye movements that transforms angular velocity inputs into angular position outputs. The necessity of such a velocity-to-position neural integrator (VPNI) was pointed out in the 1960s because sensory and command inputs to the oculomotor system encode eye velocity, whereas the extraocular motor neurons additionally carry an eye position signal (Robinson, 1968). The transformation of velocity into position is “integration” in the mathematical sense of Newtonian calculus. It has been suggested that this transformation is dependent on a relatively high degree of recurrent interactions within the VPNI. We note that the prevalence of recurrent interactions in modO was significantly higher than in modA (Figure 4A).

**Figure 4:**
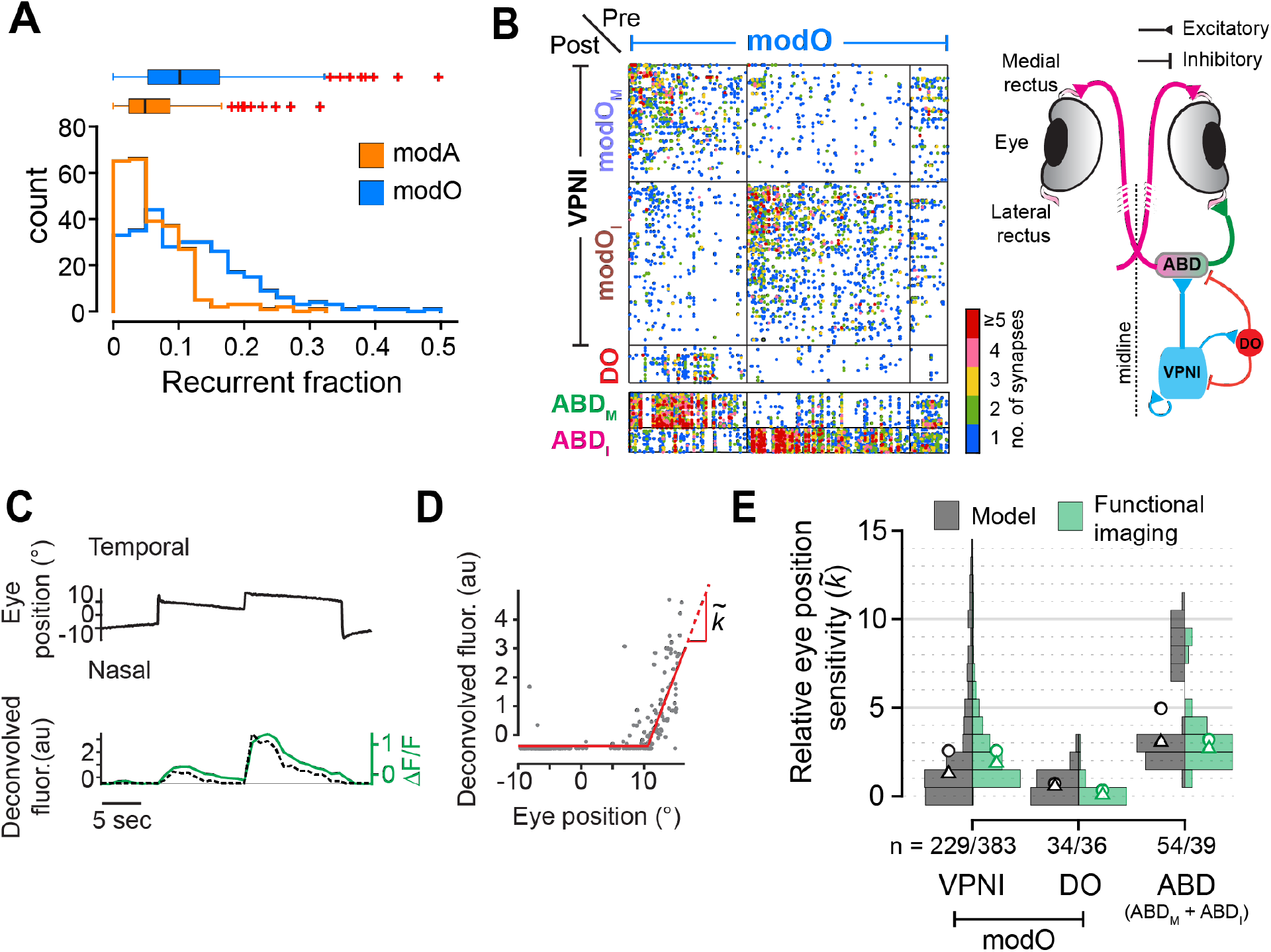
Connectome-based prediction of functional properties in the oculomotor system. **A.** Histogram of the fraction of all neurons that are recurrent within the identified modules. Box and whisker plots show medians (black line), 25^th^ and 75^th^ percentiles (box edges), and outliers (red +). **B.** (Left) Connectivity of neurons within modO, with DO neurons grouped separately for visualization. Connectivity from modO to ABD neurons in the periphery is plotted below. (Right) Schematic of a zebrafish summarizing connectivity between the different cell types, VPNI (modO_M_+modO_I_), DO, and ABD (ABD_M_+ABD_I_). **C.** Raw activity trace (green) from calcium imaging and the deconvolved fluorescence trace (dotted black line) of an example abducens neuron along with the eye position of the ipsilateral eye (black). **D.** For the neuron shown in (C), deconvolved fluorescence vs eye position (gray) and a best-fit relationship (red) that determines the relative eye position sensitivity 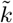. **E.** Histograms of the relative eye position sensitivity 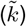 predicted from a connectome-based model (black) as compared to values determined from functional imaging experiments (green). Bimodal distribution of the ABD neurons in the model corresponds to ABD_M_ (lower) and ABD_I_ (upper) populations. Circles represent the average values and triangles represent medians. Histograms for experimental data are showing values inside the 1^st^ and 99^th^ percentile.

We developed a network model to assess if functional properties of VPNI neurons and their oculomotor partners could be predicted from connectomic information. The model consisted of four populations: DO neurons, VPNI cells, ABD interneurons, and ABD motoneurons (Figure 4B). DO neurons provide optokinetic and vestibular velocity signals to the abducens and VPNI; they exhibit little to no change in activity during spontaneous saccades and fixations (Pastor et al., 2019). VPNI neurons were identified as all neurons in modO excluding the DO cells; this was justified based on comparison of the extent of the reconstructed volume and functional maps of oculomotor cell types (see Methods). Abducens neurons receive input from both VPNI and DO neurons. We focused on predicting the relative strength of the eye position signal in these different populations during spontaneous fixation behavior.

A minimal set of physiological constraints were imposed on the model. The signs of the weights were chosen according to rationales described in the Methods. Briefly, all DO cells were inhibitory (Figure S3C, Methods) (Pastor et al., 2019), and the remaining cells were excitatory (Lee et al., 2015) (Figure S9C, Methods). The recurrent network model omitted interactions with neurons on the other side of the brain, based on previous physiological studies indicating that persistent activity in the oculomotor integrator can be generated independently by each half of the brain (Aksay et al., 2007a; Fisher et al., 2013; Gonçalves et al., 2014). Likewise, we omitted interactions between modO and modA, based upon their separate roles in controlling eye movement versus body movement behaviors (Orger et al., 2008; Huang et al., 2013; Pujala and Koyama, 2019) and the body-fixed nature of the oculomotor behavior being investigated.

To create a network model we first derived a synaptic weight matrix from our EM reconstruction as follows. Each element *W_i_j* of the weight matrix is the physiological strength of the connection received by neuron *i* from neuron *j*. We approximated *W_ij_* as proportional to *N_ij_*, the number of synapses received by *i* from *j.* This approximation effectively assumes that all synapses involved in a connection have the same strength. Then, for each neuron *i*, we divided by ∑_*k*_*N_ik_*, where the sum includes all synapses received by neuron *i*, including those from neurons outside modO. Therefore the final synaptic weight matrix took the form 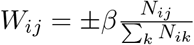.

The normalizing factor in the denominator has two interpretations. Scaling the strength of an individual synapse by the total number of synapses incoming to a neuron is similar in concept to homeostatic synaptic scaling as observed experimentally (Turrigiano, 2012). Alternatively, the normalization can be viewed as a way of compensating for truncation of dendritic arbors by the borders of the EM volume. Due to the normalization, each recurrent connection strength *W_ij_* is proportional to the fraction of the total inputs that neuron j provides to neuron *i*, and the sum ∑_*j*_*W_ij_* over neurons in modO gives the “recurrent fraction” of input to neuron i in modO, defined as the fraction of synapses received from other modO neurons (Figure 4A. Average recurrent fraction within modA and modO are μ_modA_ = 0.06 ± 0.05, μ_modO_ = 0.11 ± 0.08, p = 1.2×10^-18^, Wilcoxon-rank sum test).

Linear model neurons (Cannon et al., 1983) were used for simplicity because most VPNI cells have firing rates that vary linearly with eye position for ipsilateral eye positions (McFarland and Fuchs, 1992; Aksay et al., 2000). For each neuron, the slope of this linear rate-position relationship is known as the neuron’s eye position sensitivity. At this point, we could have simulated the dynamics of the network to extract predictions of relative eye position sensitivities. This was unnecessary because the results of simulation can be derived analytically from an eigenanalysis of the synaptic weight matrix, following previous theoretical work (Seung, 1996). The eigenanalysis yielded predictions of the eye position sensitivities of DO, VPNI and downstream ABD neurons, up to an overall scale factor (Methods). Each of the four populations displayed a characteristic distribution of eye position sensitivities (Figure 4E).

To test the model prediction, we extracted eye position sensitivities from calcium imaging experiments conducted in other larval zebrafish (Methods). We assessed eye position sensitivity during spontaneous fixations in the dark (Figure 4C, D), when persistent firing depends strictly on the internal dynamics of the VPNI. Because the imaged neurons come from other individuals, they cannot be placed in one-to-one correspondence with neurons in our EM volume. Therefore we resorted to a population-level comparison. The distribution of eye position sensitivities for each population matched the model predictions quite well (Figure 4E). We only considered three classes of neurons (VPNI, DO, and ABD) in the experiments, which could not easily distinguish between ABD_M_ and ABD_I_ neurons. However, the bimodal distribution for the ABD neurons in the experiments corresponds well to a mixture of the model ABD_M_ and ABD_I_ distributions.

Remarkably, the match between model predictions and experiments in Figure 4E involved fitting of only one free parameter. As mentioned above, the overall scale of model eye position sensitivities is arbitrary, and is related to the “read-out” of eye position from neural activity. This overall scale factor was adjusted to match model with experiment for the means of the VPNI distributions. The good match between the shapes of the VPNI distributions does not depend on any further parameter fitting. The same is true for the matches between the model and experimental distributions for the other neuron classes. There is one other adjustable parameter in the model, a scale factor in the weight matrix definition that was tuned to make the network operate as an integrator, but this scale factor is not otherwise involved in the prediction of eye position sensitivities.

To test the robustness of our result, we changed the model by adjusting the threshold for dividing nodes into center and periphery (Methods). Even if the number of neurons in the center was reduced by up to 50%, the model distributions of eye position sensitivities remained similar (Figure S10A). We also corrupted our reconstructed wiring diagram by simulating errors in the automated synapse detection, and found that the population distributions of eye position sensitivities remained very similar (Figure S10C).

We wondered whether potential connectivity would have been sufficient for our network modeling. We estimated *N_ij_* by the number of potential synapses onto neuron *i* from neuron *j*, and then computed the weight matrix *W_ij_* by the normalization described above (Methods). When our weight matrix based on potential synapses was substituted into our network model, we obtained population distributions for eye position sensitivities that were qualitatively different from experimental measurements (Figure S10D).

## Discussion

Brain modules have previously been found by analyzing networks in which nodes represent brain regions (Zingg et al., 2014; Oh et al., 2014; Bota et al., 2015). We have instead considered networks in which nodes represent individual neurons, and links represent synaptic connections. The modules in our brainstem wiring diagram were validated using additional biological information, which also enabled plausible assignments of biological functions. A related study in visual cortex (Lee et al., 2016) did not yield modules that could be biologically validated or assigned functions. We proposed a hierarchical division (Figure 2, 3), but flat clustering yields similar modules (Figure S5).

When defined at cellular resolution, a module might sound similar to a neuronal cell type (Seung and Sümbül, 2014). The concepts have some relation but are different. The neurons of a module are strongly connected with each other by definition, whereas the neurons of a cell type might not be synaptically connected with each other at all.

We defined a potential synapse as a presynapse-postsynapse pair within some threshold distance, while the conventional definition is as an axodendritic apposition (Stepanyants and Chklovskii, 2005; Reimann et al., 2015). Our definition is convenient for EM, which reveals the locations of presynapses and postsynapses, but is also relevant for light microscopy if these synaptic structures are labeled. We found that some aspects of modular organization (Figs. 2D, 3C) and forward modeling of eye position sensitivities (Figure S10d) could be predicted from potential synapses if the threshold distance were 2 μm or less. Based on this number, one might jump to the conclusion that at least some of our findings could have been obtained by diffraction-limited light microscopy. However, this technique usually requires the pooling of sparsely labeled neurons reconstructed from many animals (Shepherd et al., 2005; Jefferis et al., 2007), leading to two complications. First, spatial smoothing of arbors, typically over tens of microns, may be required to dampen random fluctuations in the number of potential synapses (Shepherd et al., 2005; Stepanyants and Chklovskii, 2005). Second, registration of multiple brains introduces additional positional uncertainty. Therefore it seems challenging to replace EM by diffraction-limited light microscopy, given that our model predictions become very poor or collapse when the threshold distance for potential synapses is 10μm (Figs. 2D, 3C, Figure S10D). Novel light microscopies that beat the diffraction limit (Igarashi et al., 2018) could work in principle, but in practice have not so far yielded comparably rich information about neural circuits.

We identified modO as containing the velocity-to-position neural integrator (VPNI) of the oculomotor system. Previous physiological studies of VPNI cells mainly focused on R7/8 (Miri et al., 2011b; Lee et al., 2015; Daie et al., 2015; Vishwanathan et al., 2017). Our map of modO (Figure 2C) suggests that the VPNI should also include R4-6. The extension is consistent with a previous observation that VPNI function was only partially abolished by sizable inactivation of R7/8 (Miri et al., 2011b), and with previous reports of eye position signals in R4-6 neurons that are not abducens neurons (Ramirez and Aksay, 2018; Brysch et al., 2019).

The VPNI has served as a model system for understanding persistent neural firing (Major and Tank, 2004; Joshua and Lisberger, 2015). The VPNI also exhibits low-dimensional brain dynamics, which has been found to underlie a wide array of motor (Cannon and Robinson, 1987; Seung, 1996; Aksay et al., 2001), navigational (Blair and Sharp, 1995; Samsonovich and McNaughton, 1997; Burak and Fiete, 2006, 2009; Yoon et al., 2013; Kim et al., 2017; Turner-Evans et al., 2020) and cognitive functions (Romo et al., 1999; Miller et al., 2005; Daie et al., 2015; Inagaki et al., 2019). Twenty-five years ago, “line attractor” and “ring attractor” network models were proposed for low-dimensional neural dynamics in the oculomotor (Seung, 1996) and head direction systems (Skaggs et al., 1995; Zhang, 1996), respectively. Connectomic information from the *Drosophila* head direction system (Turner-Evans et al., 2020) is currently being used to inform ring attractor network models (Skaggs et al., 1995; Zhang, 1996). Our work similarly constrains attractor network models of the oculomotor system (Seung, 1996; Fisher et al., 2013). This is the beginning of a trend in which connectomics will aid network modeling of low-dimensional neural dynamics, a general phenomenon that has been recognized in many brain regions and species (Yoon et al., 2013; Daie et al., 2015; Kim et al., 2017; Green et al., 2017; Vyas et al., 2020).

In most neural network models of brain function, the synaptic weight matrix has been regarded as the solution to an “inverse problem.” Given the observed effects (neural activity and behavior), the modeler attempts to identify the unobserved cause (weight matrix). Connectomics offers the possibility of treating network modeling as more of a “forward problem.” The forward approach has been feasible for small nervous systems, in which the weight matrix can be directly observed and completely mapped by synaptic physiology (Hartline, 1979).

A similar “forward” approach has been applied in dedicated sensory circuits. Wanner and Friedrich (Wanner and Friedrich, 2020) demonstrated how a connectome could be used to model whitening of odor representations in a vertebrate olfactory bulb (Wanner and Friedrich, 2020). An EM wiring diagram was used to constrain a model of orientation and direction selectivity in the *Drosophila* visual motion detection circuit (Tschopp et al., 2018), though fine-tuning by backpropagation learning was necessary. At a lower, “mesoscopic” level of resolution, inter-area projection maps in primates have been used to explain the temporal dynamics of cortical responses (Wang et al., 2020).

Our forward approach started from a synaptic weight matrix estimated from EM reconstruction, and succeeded in predicting the statistical distribution of relative eye position sensitivities for several neural populations measured by calcium imaging of animals during ocular fixations (Figure 4C). Some “inverse” aspects to our modeling remained, because we constrained ourselves to modeling eye movements in the absence of body movements and used known signs of connections (Lee et al., 2015; Pastor et al., 2019), physiological observations about the approximate linearity of oculomotor responses (Aksay et al., 2000, 2001) and independence of the bilateral halves of the circuit (Aksay et al., 2007b; Debowy and Baker, 2011).

Our success in modeling eye position sensitivities through a forward approach with minimal physiological constraints is perhaps surprising. This is especially so given our naive estimates of synaptic weights from the simple measure of number of synapses, simplified linear rate model neuron treatment of cell morphology, synaptic and intrinsic cellular biophysics, and our neglect of neuromodulation (Bargmann and Marder, 2013). Comparisons between model predictions and physiological data at the level of single cells, rather than populations, might require more sophisticated modeling of cell- and synapse-specific biophysics.

As more connectomes become available in other settings, it will be important to consider which physiological constraints need to be incorporated to make appropriate use of these powerful datasets. Answers to this question depend on many factors including the breadth of behaviors to be produced in a single model, the range of dynamics of the constituent components, and the degree to which the model is to produce quantitative versus qualitative matches to data. Nevertheless, we hope our work suggests how, when guided by knowledge of the various behaviors a circuit participates in, and appropriate physiological constraints gleaned from recordings and perturbations of activity, it may be possible in even more complex circuits to identify physiological modules whose function can be well understood using the connectome-based analysis and modeling approach taken here.

## Author Contributions

AV - Designed and conceptualized study, collected data, data interpretation, imaged EM data, performed registration, data analysis and wrote paper. ADR - collected light microscopic data, registered and analyzed LM data, data interpretation. JW - code for data generation and curation in eyewire, meshing, skeletonization, convnet inference. AS- computational modeling, RY - clustering algorithm comparisons, NK - Eyewire data assembly. DI - EM image assembly. NT - Synapses detection and partner assignment. KL - Boundary detection. IT - Eyewire algorithms development. WMS- Eyewire algorithms development, Data manipulation software. CSJ - Eyewire algorithms development, Eyewire system administration. CD, DB - Eyewire moderation, data curation. MSG - designed and conceptualized computational model, data interpretation, wrote the paper. ERFA - designed and conceptualized study, data interpretation, wrote the paper. HSS- designed and conceptualized study, data analysis, data interpretation, wrote the paper. EyeWirers - Neuron reconstruction online.

## Acknowledgement

We thank Merlin Moore, Kyle Wille, Ryan Willie, Selden Koolman, Sarah Morejohn, Ben Silverman, Doug Bland, Celia David, Sujata Reddy, Anthony Pelegrino, Sarah Williams and Dan Visser for manual annotation and proof-reading of neuron reconstructions, Amy Sterling for EyeWire management. We thank Will Wong for help with image data transformation for Eyewire and Alex Bae for help with skeletonization. We also thank Misha Tsodyks, David Kleinfeld, Carlos Brody, Misha Ahrens, Minoru Koyama, Aristides Arrenberg, Christian Brysch, Kayvon Daie, Vishwas Mishra, and Chanwoo Chun for their suggestions. ERFA, MSG, AV and HSS acknowledge support from R01 NS104926, R01 EY027036. ERFA and MSG acknowledge support from R01 EY021581, Simons Foundation Global Brain Initiative. MSG acknowledges support from the NIH-NINDS Brain initiative award 5U19NS10468-2. ADR received support from K99 EY027017. HSS acknowledges support from NIH/NCI UH2 CA203710, ARO W911NF-12-1-0594, and the Mathers Foundation, as well as assistance from Google, Amazon, and Intel. HSS is grateful for support from the Intelligence Advanced Research Projects Activity (IARPA) via Department of Interior/Interior Business Center (DoI/IBC) contract number D16PC0005. The U.S. Government is authorized to reproduce and distribute reprints for Governmental purposes notwithstanding any copyright annotation thereon.

## Methods

### Image acquisition and alignment

We acquired a dataset of the larval zebrafish hindbrain that extended 250 μm rostrocaudally and includes rhombomeres 4 through 7/8 (R4 to R7/8). The volume extends 120 μm laterally from the midline and 80 μm ventrally from the plane of the Mauthner cell axon. The ssEM dataset was an extension of the original dataset in ref Vishwanathan et al. (2017) and was extended by additional imaging of the same serial sections. Only a few tens of neurons had been manually reconstructed in our original publication on the ssEM dataset Vishwanathan et al. (2017). The dataset was stitched and aligned using a custom package, Alembic (see Code availability). The tiles from each section were first montaged in 2D, and then registered and aligned in 3D as whole sections. Point correspondences were generated by block matching via normalized cross-correlations both between tiles and across sections. The final set of parameters that were used are listed in table.

**Table1:** Parameters used for image alignment

| Step | Images | BlockMatches | Transformation | Typical block radius (pixels) | Typical search radius (pixels) | Scale | Bandpass (after downsampling) |
| --- | --- | --- | --- | --- | --- | --- | --- |
| Premontage | Tiles (8k x 8k) | 1 between each pair of adjacent tiles | Translation | Entire overlap | Entire overlap | 0.5 | (0,10) |
| Elastic Montage | Tiles (8k x 8k) | Regular triangular mesh, 100 px, in the overlapping regions between adjacent tiles | Elastic | 120 | 150 | 0.5 | (0,20) |
| Pre-Prealignment | sections | 1 between each adjacent section | Translation | 35% of images | entire images | 0.03 |  |
| Prealignment | sections | Regular triangular mesh, 2000 px, between sections | Regularized affine | 800 | 3500 | 0.25 | (2.5,12.5) |
| Rough alignment | sections | Regular triangular mesh, 375 px, between sections | Elastic | 500 | 300 | 0.25 | (2.5,10) |
| Fine alignment | sections | Regular triangular mesh, 100 px, between section | Elastic | 300 | 70 | 1 | (2.5,15) |

Errors in each step were found by a combination of programmed flags (such as lower than expected correspondences, small search radius, large distribution of norms, or high residuals after mesh relaxation) and visual inspection. They were corrected by either changing the parameters or by manual filtering of points. In most cases, the template and the source were both passed through a band-pass filter. Stitching of tiles (montaging) within a single section was split into a linear translation step (premontage) and a non-linear elastic step (elastic montage). In the premontage step individual tiles were assembled to have 10% overlap between neighboring tiles, as specified during imaging, and by fixing a single tile (anchoring) in place. They were then translated row by row and column by column according to the single correspondence found between the overlaps. In the elastic montage step, the locations of the tiles were initialized from the translated locations found previously, and blockmatches were computed every 100 pixels on a regular triangular mesh (see Table for parameters used).

Once the correspondences were found, outliers were filtered by checking the spatial distribution of the cross-correlogram (sigma filter), height of the peak of the correlogram (r value), dynamic range of the source patch contrast, kurtosis of the source patch, local consensus (average of immediate neighbors), and global consensus (inside the section). After the errors had been corrected, by filtering bad matches, the linear system was solved using conjugate gradient descent. The mean residual errors were in the range of 0.5 - 1.0 pixels after relaxation. The inter-section alignment was split into a translation step (pre-prealignment), a regularized affine step (prealignment), a fast coarse elastic step (rough alignment), and a slow fine elastic step (fine alignment). In the pre-prealignment step, a central patch of the given montaged section was matched to the previous montaged section to obtain the rough translation between two montaged sections. In the prealignment step, the montaged images were offset by that translation, and then a small number of correspondences were found between the two montaged sections, which were solved for a least-squared-residual affine transform, regularized with 10% (empirically derived) of identity transformation to reduce shear from propagating across multiple sections. Proceeding sequentially allowed the entire stack to get roughly in place. The mean residual errors were in the range of 3.5 pixels after relaxation.

### Convolutional Net Training

#### Dataset

Four expert brain image analysts (Daan Visser, Kyle Willie, Merlin Moore, and Selden Koolman) manually segmented neuronal cell boundaries from six subvolumes of EM images with VAST (Berger et al., 2018), labeling 194.4 million voxels in total. These labeled subvolumes were used as the ground truth for training convolutional networks to detect neuronal boundaries. We used 187.7 million voxels for training and reserved 6.7 million voxels for validation.

#### Network architecture

To detect neuronal boundaries, we used a multiscale 3D convolutional network architecture similar to the boundary detector in (Zung et al., 2017). This architecture was similar to U-Net (Ronneberger et al., 2015), but with more pathways between scales. We augmented the original architecture of (Zung et al., 2017) with two modifications. First, we added a “lateral” convolution between every pair of horizontally adjacent layers (i.e. between feature maps at the same scale). Second, we used batch normalization (Ioffe and Szegedy, 2015) at every layer (except for the output layer). These two architectural modifications were found to improve boundary detection accuracy and stabilize/speed-up training, respectively. For more details, we refer the reader to the Supplementary Section A.1 and Figure S10 in (Zung et al., 2017).

#### Training procedures

We implemented the training and inference of our boundary detectors with the Caffe deep learning framework (Jia et al., 2014). We trained the networks on a single Titan X Pascal GPU. We optimized the binary cross-entropy loss with the Adam optimizer (Kingma and Ba, 2014), initialized with α = 0.001, β1 = 0.9, β2 = 0.999, and ε = 0.01. The step size α was halved when the validation loss plateaued, three times during training at 135K, 145K, and 175K iterations. We used a single training example (minibatch of size 1) to compute gradients for each training iteration. The gradient for target affinities (the degree to which image pixels are grouped together) in each training example was reweighted dynamically to compensate for the high imbalance between target classes (i.e. low and high affinities). Specifically, we weighted each affinity inversely proportional to the class frequency, which was computed independently within each of the three affinity maps (x, y, and z) and dynamically in each training example. We augmented training data using (1) random flips and rotations by 90°, (2) brightness and contrast augmentation, (3) random warping by combining five types of linear transformation (continuous rotation, shear, twist, scale and perspective stretch), and (4) misalignment augmentation (K. Lee et al. 2017) with the maximum displacement of 20 pixels in x- and y-dimension. The training was terminated after 1 million iterations, which took about two weeks. We chose the model with the lowest validation loss at 550K iterations.

#### Convolutional Net Inference

The above trained network was used to produce an affinity map of the whole dataset using the ChunkFlow.jl package (git link, (Wu et al., 2019)). Briefly, the computational tasks were defined in a JSON formatted string and submitted to a queue in Amazon Web Services Simple Queue Service (AWS SQS). We launched 13 computational workers locally with NVIDIA TitanX GPU. The workers fetched tasks from the AWS SQS queue and performed the computation. The workers first cut out a chunk of the image volume using BigArrays.jl (git link) and decompose it into overlapping patches. The patches were fed into the convolutional network model to perform inference in PyTorch (Paszke et al., 2017). The output affinity map patches were blended in a buffer chunk. The output chunk was cropped around the margin to reduce boundary effects. The final affinity map chunk, which is aligned with block size in cloud storage, was uploaded using BigArrays.jl. Both image and affinity map volumes were stored in Neuroglancer precomputed format (https://neurodata.io/help/precomputed/). The inference took about 17 days in total and produced about 26 terabytes of affinity map.

#### Chunk-wise Segmentation

In order to perform segmentation of the entire volume, we divided the volume into ‘chunks’. Overlapping affinity map chunks were cut out using BigArrays.jl, and a size-dependent watershed algorithm (Zlateski and Sebastian Seung, 2015) was applied to agglomerate neighboring voxels to make supervoxels. The agglomerated supervoxels are represented as a graph where the supervoxels are nodes, and the mean affinity values between contacting supervoxels are the edge weights. A minimum spanning tree was constructed from the graph by recursively merging the highest weight edges. This over-segmented volume containing all supervoxels and the minimum spanning tree was ingested into Eyewire (https://eyewire.org) for crowdsourced proofreading.

#### Semi-automated reconstructions on Eyewire

Neurons were chosen for proofreading in Eyewire based on an initial set of ‘seed’ neurons that were identified as carrying eye position signals, by co-registering the EM volume to calcium imaging performed on the same animal (Vishwanathan et al., 2017). All pre- and postsynaptic partners of the initial seed of 22 neurons were reconstructed. Following this we reconstructed partners of the neurons that were reconstructed in the initial round in a random manner. Eyewirers were provided the option of agglomerating (merging) supervoxels using a slider to change the threshold of agglomeration. To ensure accurate reconstructions, we did two things: (1) only players who met a certain threshold, determined by their accuracy on a previously published retinal dataset (Kim et al., 2014; Bae et al., 2018) were allowed to reconstruct zebrafish neurons and (2) the reconstructions were performed by two players in two rounds, in which the second player could modify the first player’s reconstruction (Supplementary Information). Finally, after two rounds of reconstruction, neurons were validated by expert in-house image analysts, who each have more than 5000 hrs of experience. The resulting accuracy of the players in the crowd as compared to experts (assuming experts are 100%) was >80% in the first round and ~95% after the second round of tracing. The validated reconstructions were subsequently skeletonized for analysis purposes. Player accuracy was calculated as an F1 score, where F1 = 2TP / (2TP+FP+FN), where TP represents true positives, FP represents false positives, and FN represents false negatives. All scores were calculated as a sum over voxels. TP was assigned when both the player and the expert agreed the segmentation was correct. FN was assigned when the player missed segments that were added in by the expert. FP was assigned when the player erroneously added segments that did not belong. Two F1 scores were calculated for each player, once for round 1 and once for round 2. No player played the same neuron in both rounds. Typically at an agglomeration threshold of 0.3 the segmentation had an F1 score of 62%.

#### Skeletonization

The neuron segmentation IDs were ingested to an AWS SQS queue and multiple distributed workers were launched in Google Cloud using kubernetes. Each worker fetched the segmentation chunks associated with a neuron ID. The segmentation voxels were extracted as a point cloud and the Distance to Boundary Field (DBF) was computed inside each chunk. Finally a modified version of the skeletonization algorithm TEASAR was applied (Sato et al., 2000). Briefly, we constructed a weighted undirected graph from the point cloud, where the neighboring points are connected with an edge and the edge weight is computed from the DBF. Then, we took the point with the largest DBF as source, and found the furthest point as the target. The shortest path from source to target in the graph was computed as the skeleton nodes. The surrounding points were labeled as visited, and the closest remaining unvisited point was taken as the new source. We repeated this process until all the points were visited. The skeleton node diameter was set as its DBF. The skeleton nodes were post-processed by removing redundant nodes, removing ‘hairs’, based on diameter, removing branches inside soma, downsampling the nodes, merging single-child segments, and smoothening the skeleton path. All skeletonization was performed at MIP level 4

#### Synapse detection

Synapses were automatically segmented in this dataset using neural networks to detect clefts and assign the correct partner as previously described (Turner et al., 2020). Briefly, a subset of the imaged data (219*μm*^3^) was selected for annotation. The annotations were performed using the manual annotation tool VAST (Berger et al., 2018). Trained human annotators labeled the voxels that were part of the postsynaptic density (PSD) and presynaptic docked vesicle pools. A convolutional neural network was trained to match the PSD, using 107*μm*^3^ as a training set, and 36*μm*^3^ as a validation set, leaving the remaining 76*μm*^3^ as an initial test set. All of these sets were compared to the predictions of the model tuned to an F-Score of 1.5 on the validation set in order to bias towards recall, where recall = TP / (TP + FN). Biasing the predictor towards recall reduces false negatives at the cost of more false positives, which are easier to correct. Apparent human errors were corrected, and training was restarted with a new model. We also later expanded the test set by proofreading similar automated results applied to new sections of the datasets (to increase representation of rare structures in the full image volume). The final model used a RS-UNet architecture (Lee et al., 2017) implemented using PyTorch (Paszke et al., 2017), and was trained using a manual learning rate schedule, decreasing the rate by a factor of 10 when the smoothed validation error converged. The final network reached 86% precision and 83% recall on the test set after 230k training iterations.

A convolutional network was also trained to assign synaptic partners to each predicted cleft as previously described (Turner et al., 2020). All 361 synapses in the ground truth were labeled with their synaptic partners, and the partner network used 204 synapses as a training set, 73 as a validation set, and the remaining 84 as a test set. The final network was 95% accurate in assigning the correct partners of the test set after 380k training iterations.

The final cleft network was applied across the entire image volume, and formed discrete predictions of synaptic clefts by running a distributed version of connected components. Each cleft was assigned synaptic partners by applying the partner network to each predicted cleft within non-overlapping regions of the dataset (1024 x 1024 x 1792 voxels each). In the case where a cleft spanned multiple regions, the assignment within the region that contained the most of that cleft was accepted, and the others were discarded. Cleft regions whose centroid coordinates were within 1μm and were assigned the same synaptic partners were merged together in order to merge artificially split components.

Finally, spurious synapse assignments (i.e postsynapses on axons and presynapses on dendrites) were cleaned by querying the identity of the 10 nearest synapses to every synapse, where each synapse was associated with its closest skeleton node on both the pre- and post-synaptic sides. If the majority of the 10 nearest neighbors were of the same identity (pre or post), then the synapse was assigned correctly. If the majority were of an opposing identity, these synapses were assigned wrongly and were deleted. This process eliminated 1975 falsely assigned synapses (~2% of the total).

### Registration to reference atlas

Registration of the EM dataset to the Z-Brain reference atlas (Randlett et al., 2015) was carried out in two stages. We created an intermediate EM stack from the low resolution (270 nm/pixel) EM images of the entire larval brain tissue. This intermediate stack had the advantage of a similar field of view as compared to the LM reference volume, while also being of the same imaging modality as the high-resolution EM stack. The low-resolution EM stack was registered to the reference brain by fitting an affine transform that maps the entire EM volume onto the LM volume. To do this, we selected corresponding points such as neuronal clusters and fiber tracts using the tool BigWarp (Bogovic et al., 2016). These corresponding points were used to determine an affine transform using the MATLAB least squares solver *(mldivide).* Subsequently, the intermediate EM stack, in the same reference frame as the Z-Brain atlas, was used as the template to register the high-resolution EM stack onto it. This was performed in a similar manner by selecting corresponding points and fitting an affine transform. The resulting transform would transform points from the high-resolution EM space to the reference atlas space. This transform was used to map the reconstructed skeletons from high-resolution EM space to the reference atlas space.

### Identification of ABD neurons

We identified abducens motor neurons (ABD_M_, Figure 1C, Figure S3) by their overlap with the *mnx* transgenic line (S3A). ABD_M_ axons exited R5 and R6 through the abducens (VI^th^) nerve (Figure 1C, black box) as reported previously (Vishwanathan et al., 2017). Contraversive horizontal movements of the eye are driven by the medial rectus muscle, which are innervated by motor neurons in the oculomotor nucleus, which in turn are driven by inter-nuclear neurons (ABD_I_) in the contralateral abducens complex (Figure 1C). ABD_I_ neurons were identified (Figure S3A) by their overlap with two nuclei in the *evx2* transgenic line that labels glutamatergic interneurons. The ABD_I_ neurons were just dorsal and caudal to the ABD_M_ neurons, and their axons crossed the midline (Cabrera et al., 1992).

### Identification of DO neurons

We identified a class of secondary vestibular neurons known as Descending Octavolateral (DO) neurons (Figure 1D, brown). We observed that DO cells received synapses from primary vestibular afferents. The latter were orphan axons in R4 identified as the vestibular branch of the vestibulocochlear nerve (VIII^th^ nerve) by comparison with the *isl-2* line, which labels the major cranial nerves (Figure S3B, blue axons).

### Identification of Reticulospinal neurons

The RS neurons were divided into large and small groups (Figure1H). Large RS neurons were the M, Mid2, MiM1, Mid3i and CaD neurons. Small RS neurons were RoV3, MiV1, MiV2. These were identified by their stereotypic locations (Metcalfe et al., 1986) and by comparison within the Z-Brain atlas (Figure S4).

### Centrality-based division into center and periphery

The division of the reconstructed wiring diagram into center and periphery is based on standard measures of “centrality” which have been devised in network science (Newman, 2018). We define the simplest measure, known as degree centrality, as the geometric mean of the in-degree and out-degree of a node. (It is more common to choose one or the other.) Another popular measure, known as eigenvector centrality or eigencentrality, is a node’s element in the eigenvector of the connection matrix (number of synapses onto node *i* from node *j*) corresponding to the eigenvalue with maximal real part--this measure extends the simpler concept of degree centrality by weighing a node’s connections by their centrality, i.e. a node is more central to the network if it receives inputs from other high centrality neurons. Mathematically, this defines the eigenvector problem *v* = ∑*N_ij_v_j_* where *v_i_* is the (input) centrality of neuron i. An analogous formula, but instead replacing *N_ij_* by its transpose (thus, defining left rather than right eigenvectors) can be used to weight output connections by the (output) centrality of the node to which output is projected. The eigenvector elements can be chosen non-negative by the Perron-Frobenius theorem. It is standard to use either the left or right eigenvector, but we use both for our definition by computing the geometric mean of the left and right eigenvector elements. Degree centrality and eigencentrality are correlated, but not perfectly (Figure S1C). For a visualization of the network based on eigencentrality, see Figure S1E.

Our definition of the periphery relies mainly on degree centrality; the vast majority of the periphery consists of 2282 nodes with vanishing degree centrality. We also define the periphery to include an additional 62 nodes with vanishing (<10^-8^) eigencentrality but nonzero degree centrality. The remaining 540 recurrently connected neurons are defined as the “center” of the graph. (See below for effects of varying the eigencentrality threshold for center-periphery division.)

### Graph Clustering

We applied three graph-clustering algorithms to divide the center into modules, and obtained similar results from all three. The clustering from the Louvain algorithm is presented in the main text, and those of the spectral algorithm and stochastic block model in the supplementary information.

#### Louvain Clustering

Graph clustering was performed using the Louvain clustering algorithm for identifying different ‘communities’ or ‘modules’ in an interconnected network by optimizing the ‘modularity’ of the network, where modularity measures the (weighted) density of connections within a module compared to between modules. Formally, the modularity measure maximized is *Q_gen_* = ∑*B_ij_δ*(*c_i_, c_j_*), where *δ*(*c_i_,c_j_*) equals 1 if neurons a and b are in the same module and 0 otherwise, where 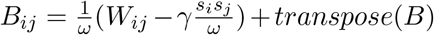. Here *s_i_* = ∑_*c*_ *W_ic_* is the sum of weights into node *i, s_j_* = ∑_*c*_ *W_jb_* is the sum of weights out of node *j*, *ω* = ∑_*cd*_ *W_cd_* is the total sum of weights in the network, and the resolution parameter *γ* determines how much the naively expected weight of connections 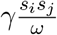 is subtracted from the connectivity matrix. Potential degeneracy in graph clustering was addressed by computing the consensus of the clustering similar to (Sporns and Betzel, 2016). Briefly, an association matrix, counting the number of times a node (neuron) is assigned to a given module, was constructed by running the Louvain algorithm 200 times. Next, a randomized association matrix was constructed by permuting the module assignment for each node. Reclustering the thresholded association matrix, where threshold was the maximum element of the randomized association matrix, provided consensus modules. We used the *commuinity_louvain.m* function from the Brain Connectivity Toolbox package (BCT, https://sites.google.com/site/bctnet/Home). In addition to the Louvain graph-clustering algorithm, we also clustered the ‘center’ with two alternate graph-clustering algorithms; spectral clustering and stochastic block matching, described below.

#### Spectral Clustering

We employed a generalized spectral clustering algorithm for weighted directed graphs to bisect the zebrafish ‘center’ subgraph as proposed by (Chung, 2005). Given a graph *G*(*V,E*) and its weighted adjacency matrix 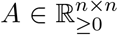, where *A_ij_* indicates the number of synapses from neuron *i* to neuron *j*, one can construct a Markov chain on the graph with a transition matrix *P_α_*, such that |*P_α_*|_*ij*_ := (1 - *α*) · *A_ij_*/∑_*k*_ *A_ik_* + *α/n*. The coefficient *α* > 0 ensures that the constructed Markov chain is irreducible, and the Perron-Frobenius theorem guarantees *P_α_* has a unique positive left eigenvector *π* with eigenvalue 1, where *π* is also called the stationary distribution. The normalized symmetric Laplacian of the Markov chain is 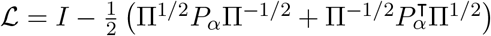.

To approximately search for the optimal cut, we utilize the Cheeger inequality for a directed graph Chung (2005) that bridges the spectral gap of 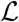 and the Cheeger constant *ϕ\**. As shown in (Gleich, 2006), the eigenvector *v* corresponding to the second smallest eigenvalue of 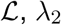, results in optimal clusters. We obtained two clusters by a binary rounding scheme, i.e., *S* = {*i* ∈ *V*|*v_i_* ≥ 0} and *S* = {*i* ∈ *V*|*v_i_* < 0}.

We modified the *directed_laplacian_matrix* function in the NetworkX package (https://networkx.github.io) to calculate the symmetric Laplacian for sparse connectivity matrices, with a default *α* = 0.05. The spectral gaps for the eigenvector-centrality subgraph is 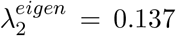 and for the partitioned oculomotor (modO) module is 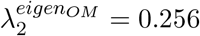.

#### Degree-corrected Stochastic Block Matching (SBM)

Unlike the Louvain and spectral clustering algorithms that assume fewer intra-cluster connections than inter-cluster connections, the stochastic block models (SBMs) do not make this assumption. We applied an efficient statistical inference algorithm (Peixoto, 2014) to obtain the SBMs that best describes the ‘center’ subgraph.

The traditional SBM (Holland et al., 1983) is composed of *n* vertices, divided into *B* blocks with {*n_r_*} vertices in each block, and with the probability, *p_rs_,* that an edge exists from block *r* to block *s*. Here we use another equivalent definition, to use average edge counts from the observation *e_rs_* = *n_r_n_s_p_rs_* to replace the probability parameters. The degree-corrected stochastic block model (Karrer and Newman, 2011) further specifies the in- and out-degree sequences 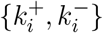 of the graph as additional parameters.

To infer the best block membership {*b_i_*} of the vertices in the observed graph *G*, we maximize likelihood 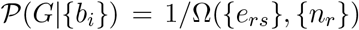, where Ω({*e_rs_*}, {*n_r_*}) is the total number of different graph realizations with the same degree distribution 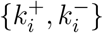, and {*e_rs_*} edges among and within blocks of sizes {*n_r_*}, corresponding to the block membership {*b_i_*}. Therefore, maximizing likelihood is equivalent to minimizing the microcanonical entropy (Bianconi, 2009) 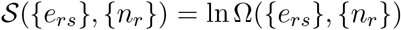, which can be calculated as 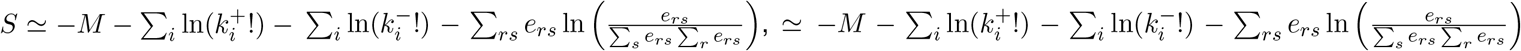, *S* where *M* = ∑_*rs*_ *e_rs_* is the total number of edges.

We used the *minimize_blockmodel_dl* function in the graph-tool package (https://graph-tool.skewed.de) to bisect the central subgraphs by setting *B_min_* = *B_max_* = 2 and *degcorr* = true.

### Potential Synapse Formalism

We define a potential synapse as a presynapse-postsynapse pair within a certain threshold distance (Figure 2D). This definition is somewhat different from the light microscopic approach, which defines a potential synapse (Stepanyants et al., 2002; Stepanyants and Chklovskii, 2005) as an approach of an axon and dendrite within some threshold distance. Also we use neurons reconstructed from a single animal, while the light microscopic approach aggregates neurons from multiple animals, or “clones” a single neuron many times Markram et al. (2015).

### Calcium imaging and eye position signals

The complete methods for recording calcium activity used to create the functional maps are reported in (Ramirez and Aksay, 2018). Briefly, we used two-photon, raster-scanning microscopy to image calcium activity from single neurons throughout the hindbrain of 7-8 day old transgenic larvae expressing nuclear-localized GCaMP6f, Tg(HuC:GCaMP6f-H2B) strain *cy73-431* from Misha Ahrens’ lab. Horizontal eye movements were recorded simultaneously with calcium signals using a substage CMOS camera. We used the CalmAn-Matlab software to extract the neuronal locations from fluorescence movies (Giovannucci et al., 2019).

We analyzed saccade-triggered average (STA) activity to determine which neurons were related to eye movements (see (Ramirez and Aksay, 2018) for complete details). For each cell, we interpolated fluorescence activity that occurred within five seconds before or after saccades to a grid of equally spaced, ⅓ second timepoints and then averaged the interpolated activity across saccades to compute the STA. Separate STAs were taken for saccades towards the left and right. We performed a one-way ANOVA on each STA to determine which neurons had significant saccade-triggered changes in average activity (p<0.01 using the Holm-Bonferroni method to correct for multiple comparisons). To determine which of these neurons had activity related to eye movement and eye velocity, we first performed a Principal Components Analysis (PCA) on the STAs from neurons with significant saccade-triggered changes. We found that the first and second principal components had post-saccadic activity largely related to eye movement and eye velocity sensitivity respectively (see Figure 3A in (Ramirez and Aksay, 2018)). We characterized each STA using a scalar index called *ϕ* in (Ramirez and Aksay, 2018), created from that STA’s projections onto the first two principal components and found that this index does a good job of characterizing the average eye movement and eye velocity-related activity seen across the population (see Figure 3C in (Ramirez and Aksay, 2018) for a map of values and population average STAs). Eye position and eye velocity neurons were defined as neurons with an STA whose value of ϕ was within a specific range (−83 to 37 and 38-68 respectively). We removed any neurons with pre-saccadic activity that was significantly correlated with time until the upcoming saccade. The locations of each neuron were then registered to the Z-Brain atlas (Randlett et al., 2015) using similar methods as listed in the previous section (see (Ramirez and Aksay, 2018) for complete details).

### Relationship between firing rate and eye position

To assess the functional characteristics of various oculomotor neurons (Figure 4) we fit a traditional model of eye position sensitivity to neuronal firing rates extracted from our fluorescence measurements. We approximated the relative firing rate, *r*, of a cell using the deconvolution algorithm with non-negativity constraint described in Pnevmatikakis et al. (2016). Comparison of relative firing rate across neurons in different populations was justified as we observed similar sensor expression levels and baseline noise levels in these populations.

We modeled the dependence of firing rate on eye position using the equation, 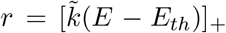, where 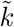 is the relative eye position sensitivity, *E_th_* is the threshold eye position at which *r* becomes positive, and the function [*x*]_+_ = *x* if *x* > 0 and 0 otherwise (Aksay et al., 2007a). In order to best compare results across animals, we normalized the units of eye position before fitting the model by subtracting the median eye position about the Null position (measured as the average raw eye position) and then dividing by the 95th percentile of the resulting positions. Since our focus was on a cell’s position-dependence, we also eliminated the eye velocity dependent burst of spiking activity at the saccade time that ABD and VPNI neurons are known to display by removing samples that occur within 1.5 seconds before or 2 seconds after each saccade. Saccade times were found in an automated fashion by determining when eye velocity crossed a threshold value (Ramirez and Aksay, 2018). Finally, since the eye position and fluorescence were recorded at different sampling rates, we linearly interpolated the values of neuronal activity at the eye position sample times.

To fit the value of 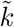, for each cell, we defined the eye movements toward the cell’s responsive direction as positive so that 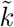 is positive by construction. We then determined the threshold, *E_th_,* using an iterative estimation technique based on a Taylor series approximation of [*x*]+ described in Ref. (Muggeo, 2003). Using the resulting estimate of *E_th_*, we determined 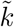 as the slope resulting from a linear regression (with offset) to *r* using *E* as a regressor. Since we do not know the cell’s responsive direction *a priori,* we ran the model twice -- once with movements to the left as positive and once with movements to the right as positive -- and used the value of 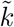 that resulted in the highest *R*^2^ value.

As a goodness-of-fit measure we required all neurons, except DO neurons, to have an *R*^2^ value greater than 0.4. Additionally, non-DO neurons were required to have a saccade-triggered average with at least one significant time point (p<0.01 by an ANOVA test using Holm-Bonferroni correction) as defined in (Ramirez and Aksay, 2018) and to have a *dF/F* response that was loosely related to eye position (*R*^2^ greater than 0.2 when we run the above model replacing *r* with *dF/F*). The relative eye position sensitivity, 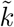, for fluorescence data was then scaled to average physiological VPNI responses from goldfish (Aksay et al., 2007a; Debowy and Baker, 2011).

### Identification of excitatory versus inhibitory neurons

VPNI neurons in the dorsomedial stripe running from R4 to R7/8 overlap in the Z-Brain atlas with a region of *alx* expression (Figure. S9C, *ZBB-alx-gal4*) where most neurons are glutamatergic (Kimura et al., 2006). More lateral and caudal neurons overlap with markers for glycine and glutamate. Finally, VPNI neurons (which have ipsilateral axonal projections) in the oculomotor module do not overlap with the GABA expression (Figure S9C, *ZBB-gad1b).* For these reasons we assigned neurons in the oculomotor module to be excitatory. DO neurons with ipsilateral projections (used in our unilateral model) have been identified as being inhibitory (Pastor et al., 2019).

### Network model based on synaptic wiring diagram

A unilateral model of the oculomotor integrator was built using the reconstructed synapses for the ABD, DO, and VPNI populations. Although the VPNI is a bilateral circuit, previous experiments (Aksay et al., 2007a; Debowy and Baker, 2011) have shown that one half of the VPNI is nevertheless capable of maintaining the ipsilateral range of eye positions after the contralateral half of the VPNI is silenced. This may reflect that most neurons in the contralateral half are below a threshold for transmitting synaptic currents to the opposite side when the eyes are at ipsilateral positions (Fisher et al., 2013). Therefore, we built a recurrent network model of one half of the VPNI circuit based on the modO neurons that we had reconstructed from one side of the zebrafish brainstem. We did not include the modA neurons, assuming that input from modA cells fell below a threshold needed to drive modO neurons, similar to the manner in which bilateral interactions have been shown to be negligible for the maintenance of persistent activity. We added the ABD neurons and the feedforward connections from modO to ABD to the model because the ABD neurons are the “read-out” of the oculomotor signals in the VPNI. Projections from the DO population were taken to be inhibitory while all other connections were taken to be excitatory, as explained above. Remaining neurons in the oculomotor module were included as part of the VPNI--we did not consider other vestibular populations since only DO neurons in modO received vestibular afferents from the VIII^th^ nerve; saccadic burst neurons and recently discovered pre-saccadic ramp neurons are likely located in R2/3 (Ramirez and Aksay, 2018), outside of our reconstructed volume.

Directed connection weights between each pair of neurons were set in proportion to the number of synapses from the presynaptic neuron onto the postsynaptic neuron divided by the total number synapses onto the postsynaptic neuron 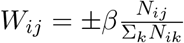. Thus, we assume that each element *W_ij_* corresponds to the fraction of total inputs to neuron *i* that are provided by neuron *j*. The scale factor *β* was set to achieve perfect integration in a linear rate model governed by 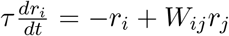, where *r_i_* is the firing rate of the *i^th^* neuron, *W_ij_* are the connection weights, and *τ*, the intrinsic time constant, is 100 ms. Fine tuning of this scale factor to achieve perfectly stable fixations is not critical to the results obtained here as the time courses of the persistent neural activity does not affect the relative firing rates of neurons that determine their relative eye position sensitivities. Position sensitivities could be determined numerically by simulating the response of the network to a pulse of input along the integrating direction. However, for a linear network, this is not necessary because the resulting relative persistent firing rates are equal to the leading eigenvector of the matrix *W_ij_* and thus can be determined analytically. Finally, the position sensitivity was multiplied by a single global scale factor that was determined by matching the average VPNI population responses from the model to the average physiological VPNI responses from goldfish (Aksay et al., 2007a; Debowy and Baker, 2011).

As a test of the robustness of our results to possible errors in connectome reconstruction, we generated a connectome that accounted for the estimated false positive and negative rate of synapse detection by our connectome reconstruction procedure. We generated 1000 models by randomly varying the identified synapses according to the estimated false positive and false negative rates and calculated the connection weights as described above. The eye position sensitivities with this synaptic detection jitter were reported as the average of these 1000 models (Figure S10C). We also tested how robust our results were to the cutoff criterion for including neurons in the recurrently connected center by progressively increasing the minimum eigenvector centrality criterion for counting a neuron as belonging to the center as opposed to periphery. We then plotted how the simulation model results changed as a function of the number of center neurons was decreased and simultaneously reported the resulting number of VPNI (i.e. non-DO neurons in modO) neurons (Figure S10A). To characterize the degradation in model performance when the actual connectome was replaced by connectomes generated by spatial proximity (potential connections), we re-ran all of the analyses described above using potential connectomes defined for connections within 2, 5, or 10μm.

### Code availability

- Alembic - https://github.com/seung-lab/Alembic.git
- BigArrays.jl - https://github.com/seung-lab/BigArrays.jl with Apache License Version 2.0.
- ChunkFlow.jl - https://github.com/seung-lab/ChunkFlow.jl with Apache License Version 2.0.
- Watershed - https://github.com/seung-lab/Watershed.jl with GNU General Public License v3.0.
- Agglomeration - https://github.com/seung-lab/Agglomeration with MIT License.
- Skeletonization, morphology and functions could be found at https://github.com/seung-lab/RealNeuralNetworks.jl with Apache License 2.0.

