## Extended figures for "Predicting modular functions and neural coding of behavior from a synaptic wiring diagram"

### Supplementary Figures

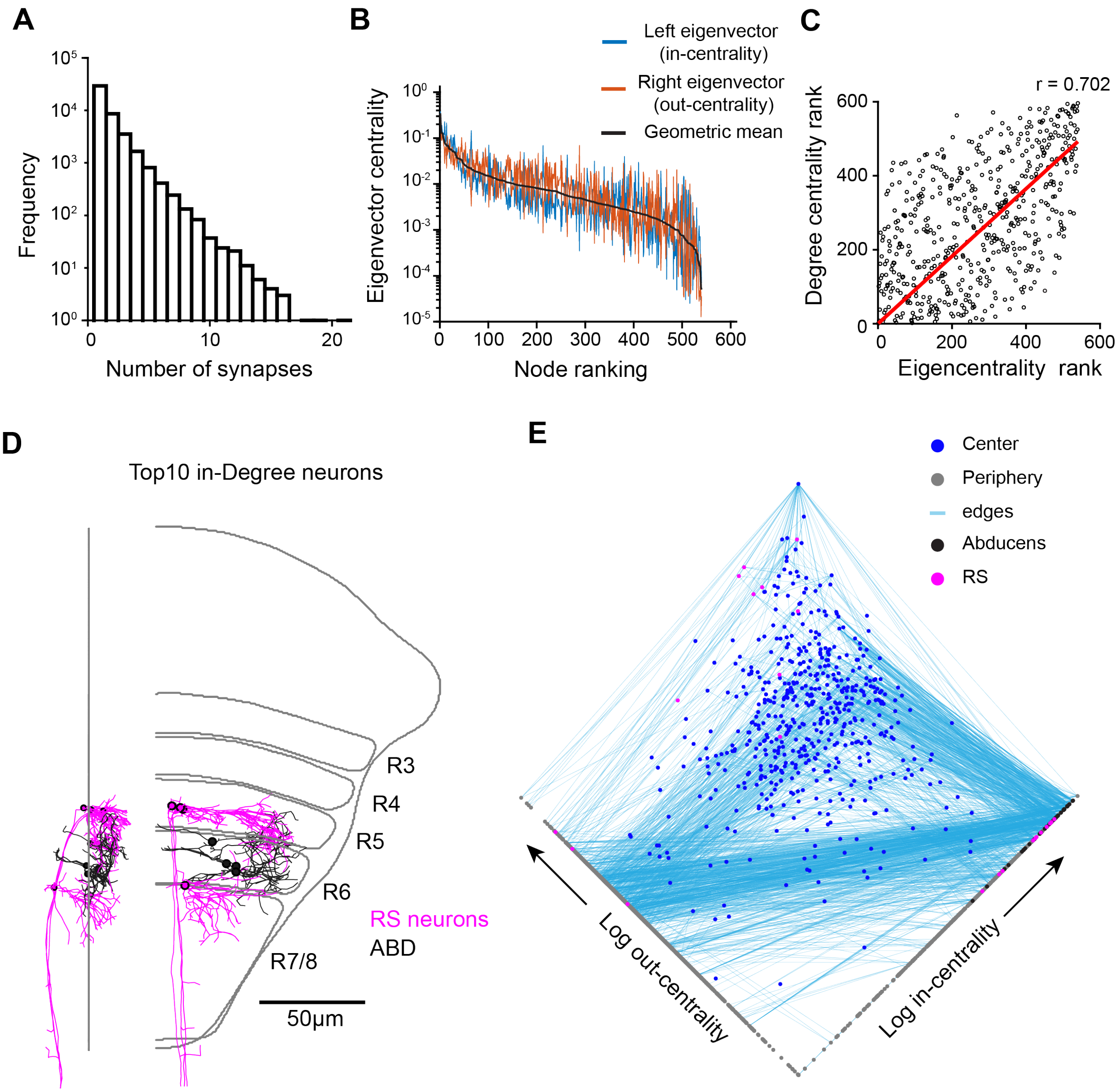


**Figure S1: Important features of the reconstructed network**

**A.** Frequency distribution of the number of synapses per connection for all neurons in the network with soma in the EM volume.

**B.** Elements of the left leading eigenvector of the connectivity matrix, the right leading eigenvector, and the geometric mean of these elements (i.e. eigencentrality) for cells ranked by eigencentrality.

**C**. The rank of neurons when ordered by degree centrality vs eigencentrality.

**D.** Top10 in-degree neurons 4 are vSPNs and 5 are ABD neurons.

**E.** Graphical representation of centrality relationships in the network. Dots represent individual neurons and lines represent the connections between neurons. Only edges involving more than 5 synapses shown for clarity.

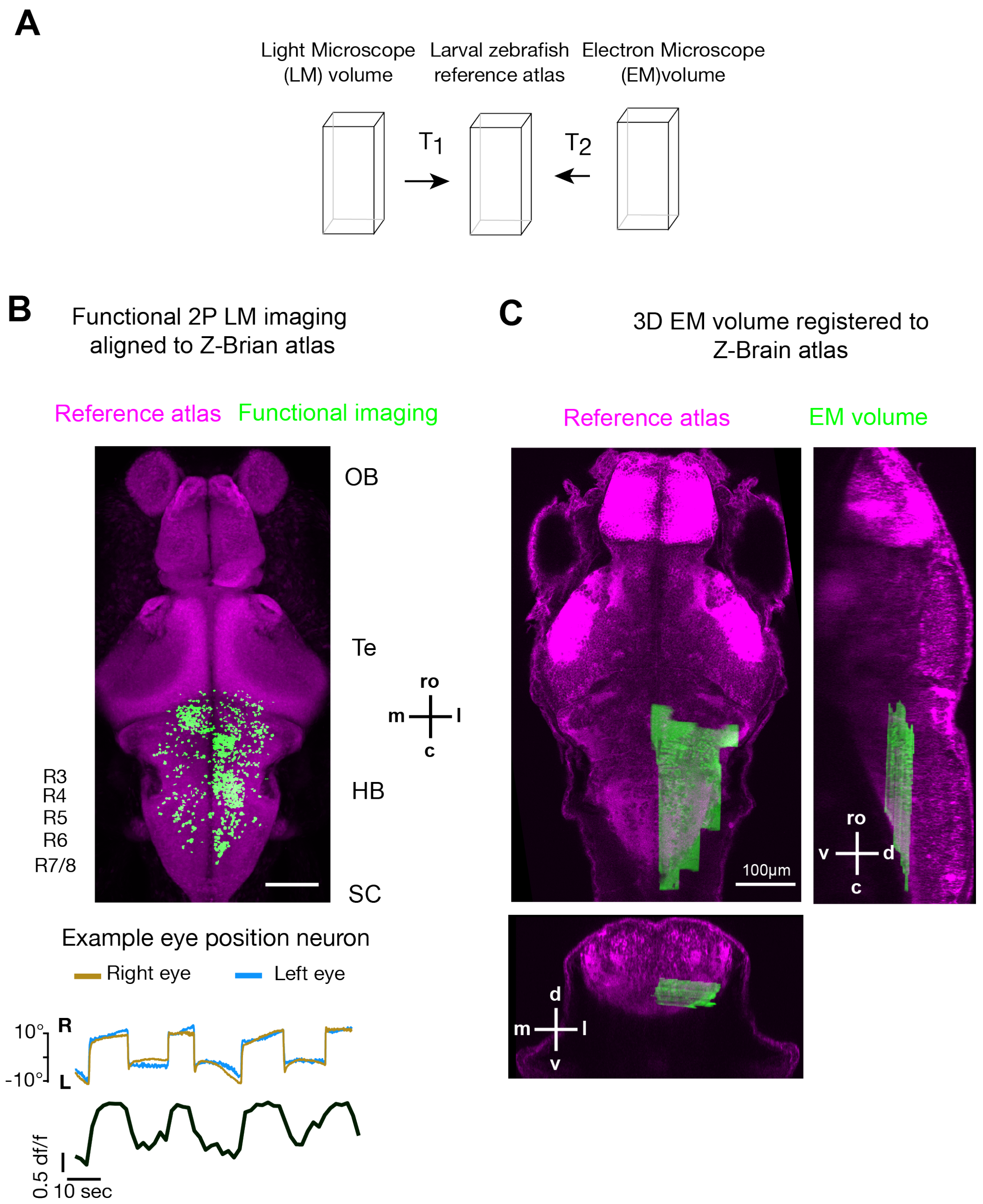


**Figure S2: Registration of electron microscopy and light microscopy images to a reference atlas**
**A.** Schematic indicated separate transformations (T) for light microscopy (LM) images and electron microscopy images onto a common zebrafish brain (Z-Brain) reference atlas.
**B.**(Top) Maximum intensity projection in the horizontal plane of registered two-photon (2P) functional imaging data (green) onto the Z-Brain reference atlas (magenta). Locations of neurons that were responsive to increase in right eye position are shown. (Bottom) Example activity of a neuron (black) during changes in eye position (blue, yellow). Scale bar 100$\mu$m.
**C.** Single plane views (top left: horizontal; top right: sagittal; bottom: coronal) of the registeredEM data set(green) onto Z-Brain reference atlas (magenta).

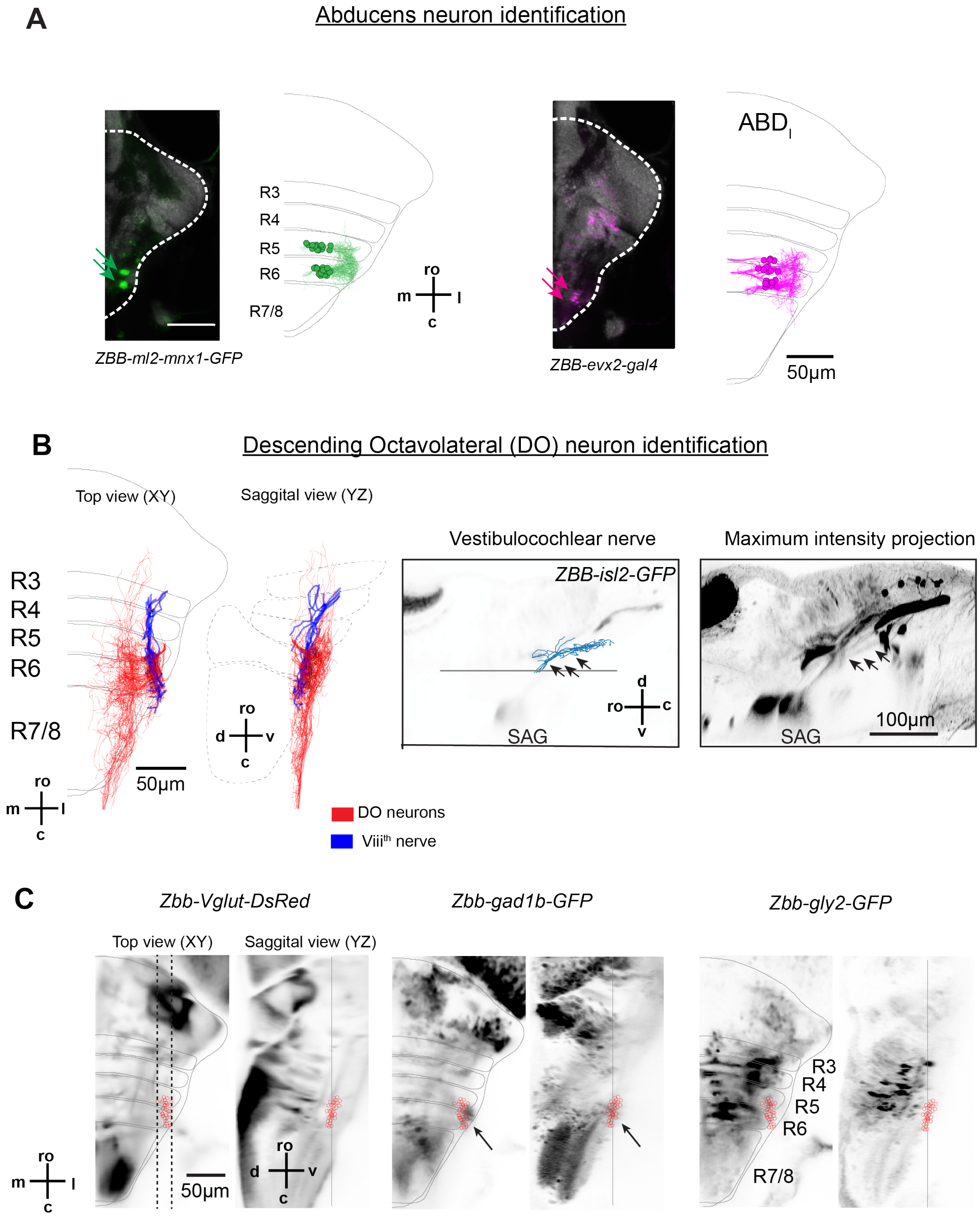


**Figure S3: Identification of abducens and descending octaval neurons.**
**A.** Overlap of abducens neurons with transgenic lines registered to the Z-Brain atlas. (Left) Location of abducens motoneurons labelled in the *ZBB-ml2-mnx1-GFP* transgenic line as indicated in fluorescence micrographs by two clusters of neurons in R5,6 (green arrows). Corresponding locations of ABDM neurons from EM reconstructions are shown in green. (Right) Location of two clusters of abducens internuclear cells labelled in the *ZBB-evx2-gal4* transgenic cross (magenta arrows) corresponds with EM reconstructions of ABDI neurons. (ro - rostral; c - causal; m - medial; l - lateral). Scale bar 100$\mu$m.
**B.** (Left) Reconstruction of descending octaval (DO) neurons (red) along with inputs (blue) from the vestibulocochlear nerve (VIII$^{th}$ nerve). Side view highlights the ventral exit of the VIII$^{th}$ nerve in R3 (arrow). Red circles indicate somata, and grey line indicates the mean horizontal plane through the DO soma. (Middle) Single plane (right) maximum intensity projection. SAG - Statoacoustic Ganglia. (ro - rostral; c - causal; d - dorsal; v - ventral). Scale bar 50$\mu$m
**C.** Overlap of DO neuron soma (red circles) with transgenic lines that indicate the location of excitatory (*ZBB-Vglut-Dsred)* and inhibitory (*ZBB-gad1b-GFP, ZBB-gly2-GFP*) neurons. The horizontal planes (top view) are the average intensity plane over the rage of DO soma along the dorso-ventral axis. Sagittal view are maximal intensity projections of the planes that contain DO soma (dotted lines). Black arrows indicate the overlap of DO neurons to a GABAergic cluster of neurons.

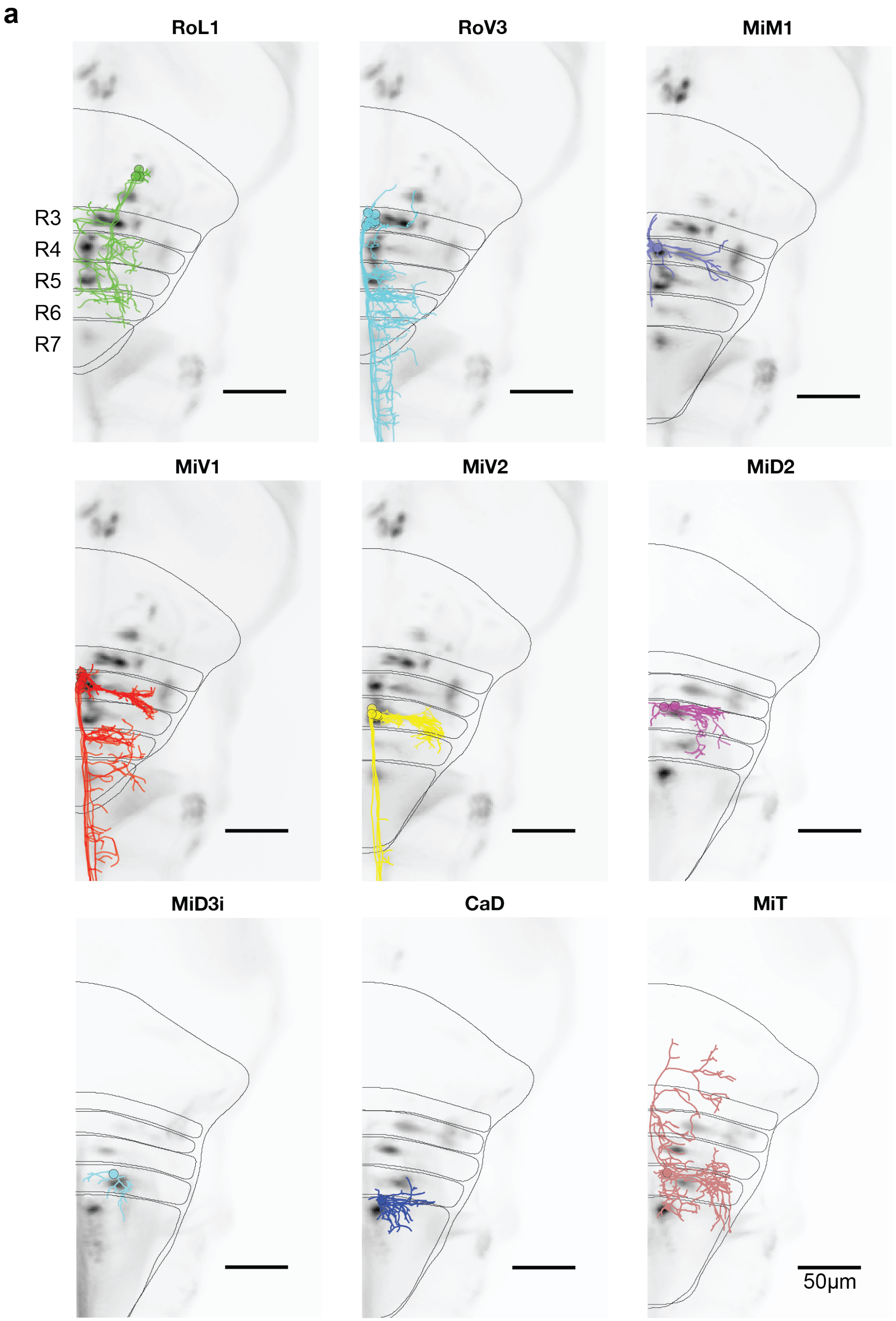


**Figure S4: Identification of vSPNs.**
**A.** vSPNs were identified based on their overlap with neurons labeled by spinal backfills that were part of the Z-Brain atlas (ro - rostral; c - causal; m - medial; l - lateral). Scale bar 50$\mu$m.

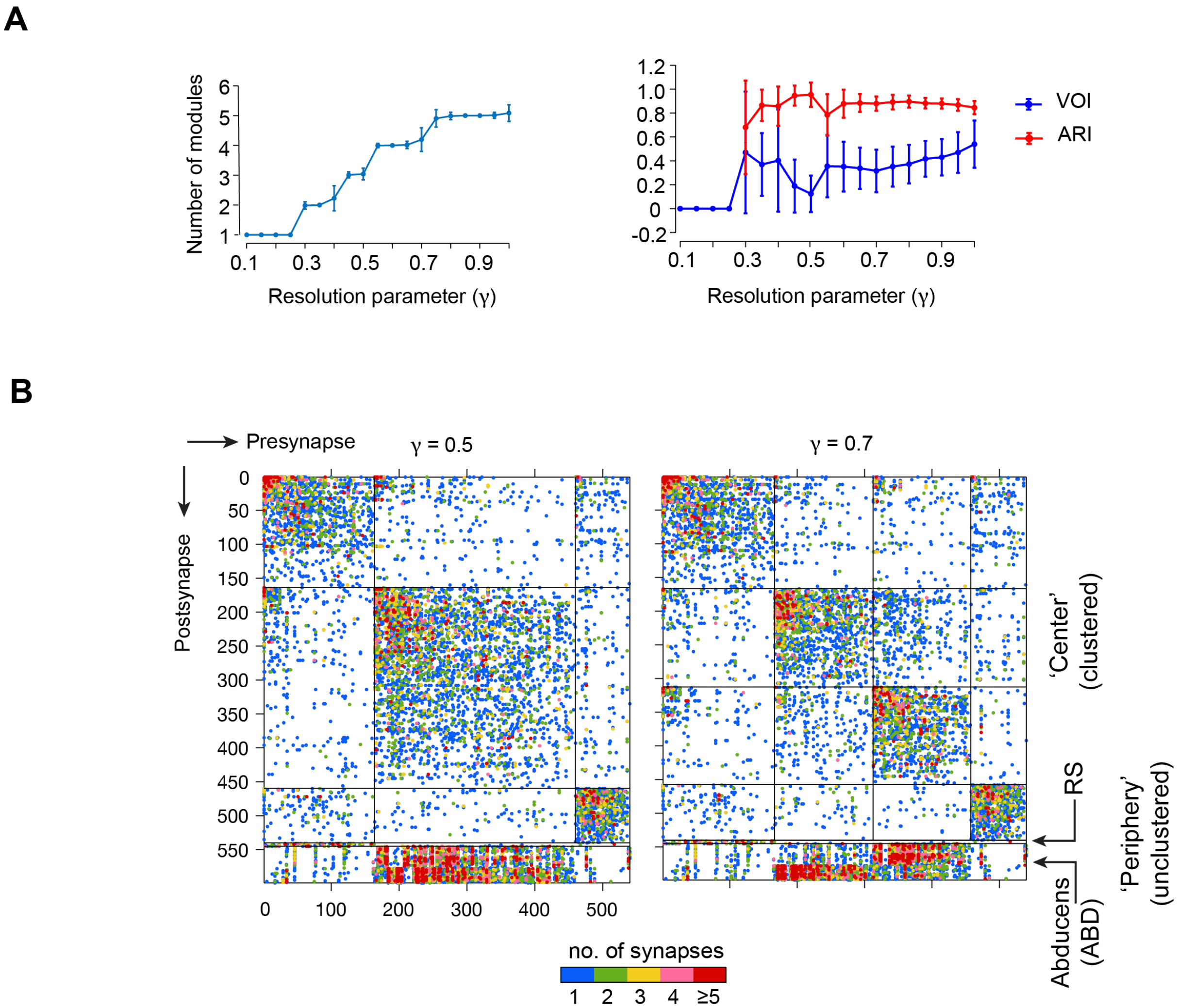


**Figure S5: Modular structure at different levels of resolution.**

**A.** Number of modules obtained and robustness of clusters as a function of resolution parameter $\gamma$.

**B.** Connectivity matrices when clustered for different numbers of modules. At $\gamma$ = 0.7, emergent modular structure is similar to that as shown when modules in Fig 2 (modA, modO) and Fig 3 ($modO_M$, $modO_I$) are clustered in a hierarchical manner.

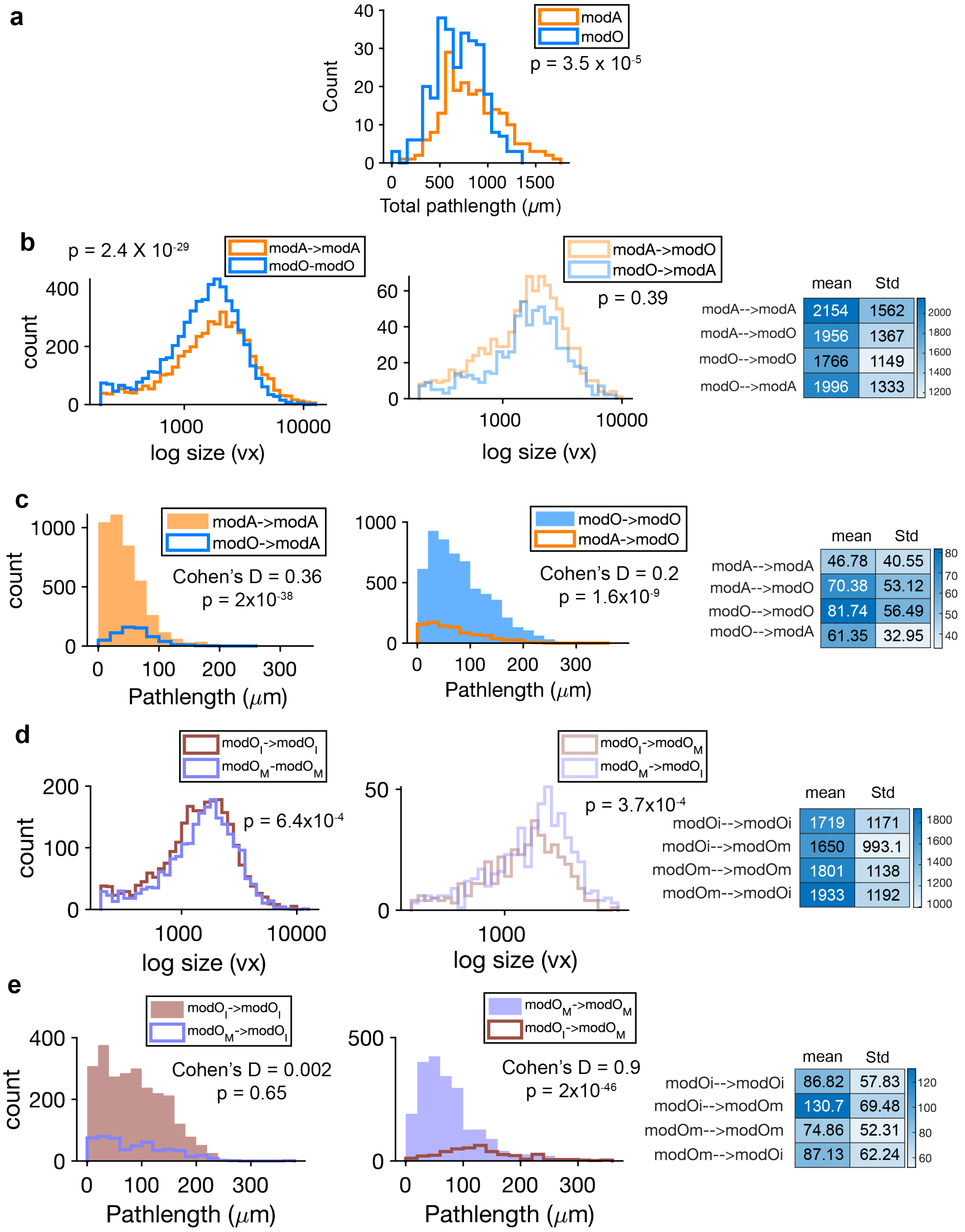


**Figure S6: Synapse size and location distribution**.
**A.** Histogram of total pathlength of neurons in modA (n = 251) and modO (n = 289). P is the significance value based on Wilcoxon-rank sum test (RS- test). Cohen's D measures the effect size.
**B.** Histogram of synaptic size (detected PSD voxels [vx]) for connections within(left) and between (right) modules modA and modO.. Table (here and below) summarizes the mean and standard deviations.
**C.** Histogram of synapses locations i.e. distance from somata to synaptic site along the neurite for neurons in modA and modO. Table summarizes the mean and standard deviations.
**D.** Histogram of synapses size (detected PSD) of neurons within-module (left) and between-modules (right) for $modO_I$ and $modO_M$. Table summarizes the mean and standard deviations.
**E.** Histogram of synapse locations for neurons within the oculomotor module, $modO_I$ and $modO_M$.

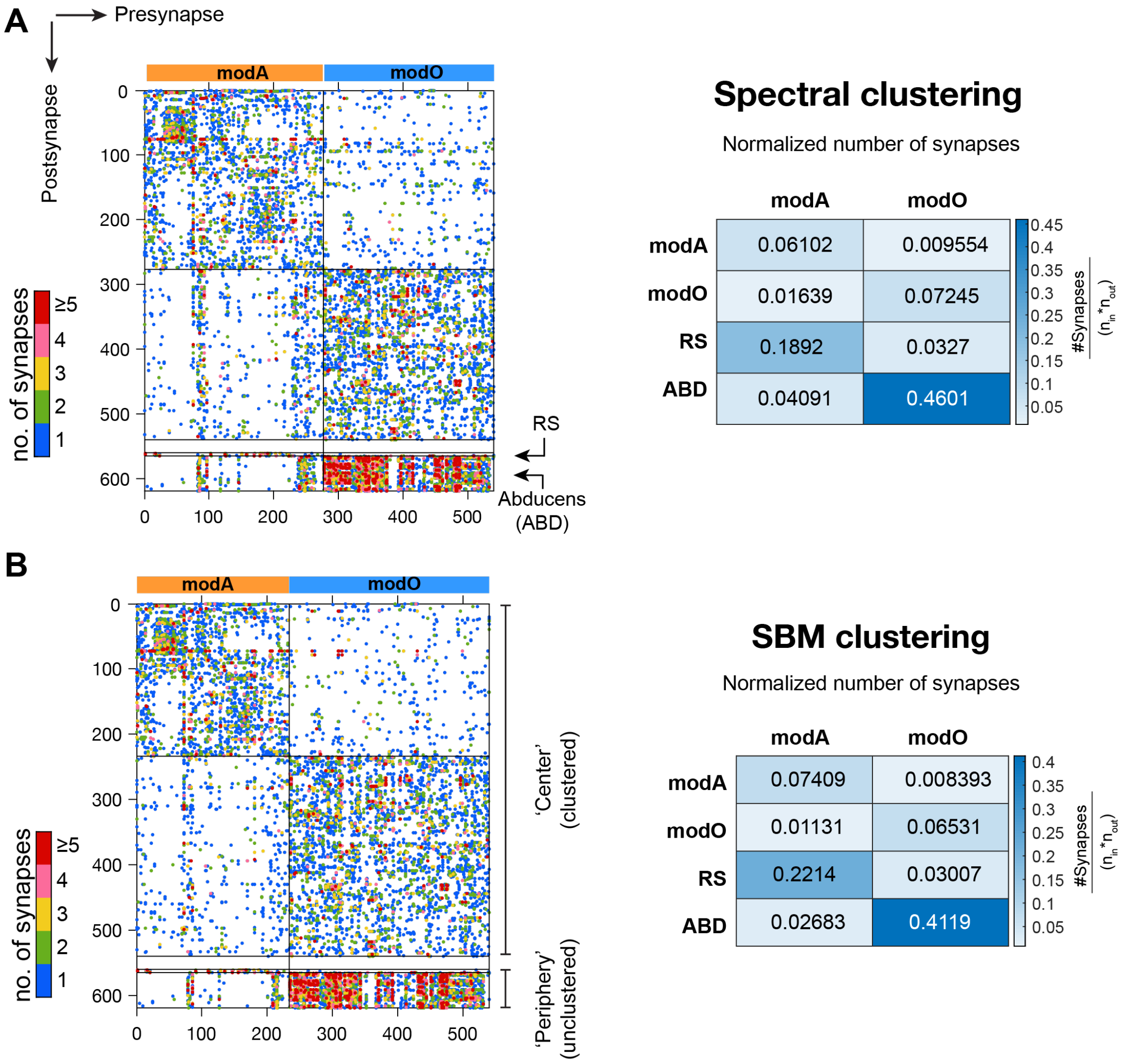


**Figure S7: Clustering algorithm comparisons**.

**A.** (Left) Connectivity when the center is organized by spectral clustering into two modules. (Right) Normalized number of synapses within and between relevant cell groups.

**B.** (Left) Connectivity when the center is organized by degree corrected Stochastic Block Matching (SBM) into two modules. (Right) Normalized number of synapses within and between relevant cell groups.

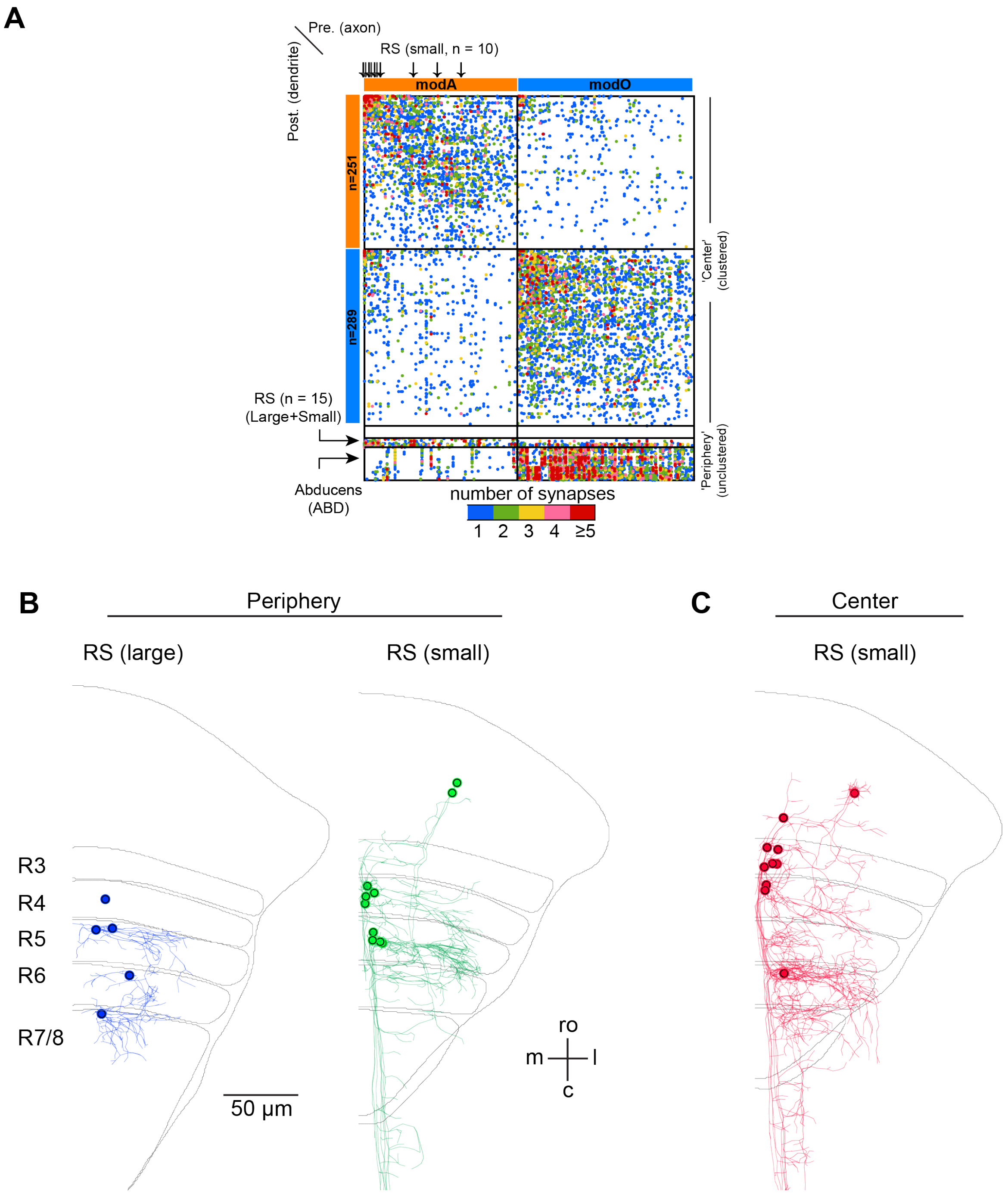


**Figure S8: Composition of Axial module (modA).**
**A.** Connectivity matrix (as in Figure. 2) with identification (arrows) of small RS neurons that were part of the center and inclusion into the periphery the remaining small RS neurons along with the large RS neurons.
**B.** Visualization of the large and small classes of RS neurons in the periphery (ro - rostral; c - causal; m - medial; l - lateral).
**C.** Visualization of small vSPNs that are part of the 'center' in modA and are indicated by arrows above the rows in (A).

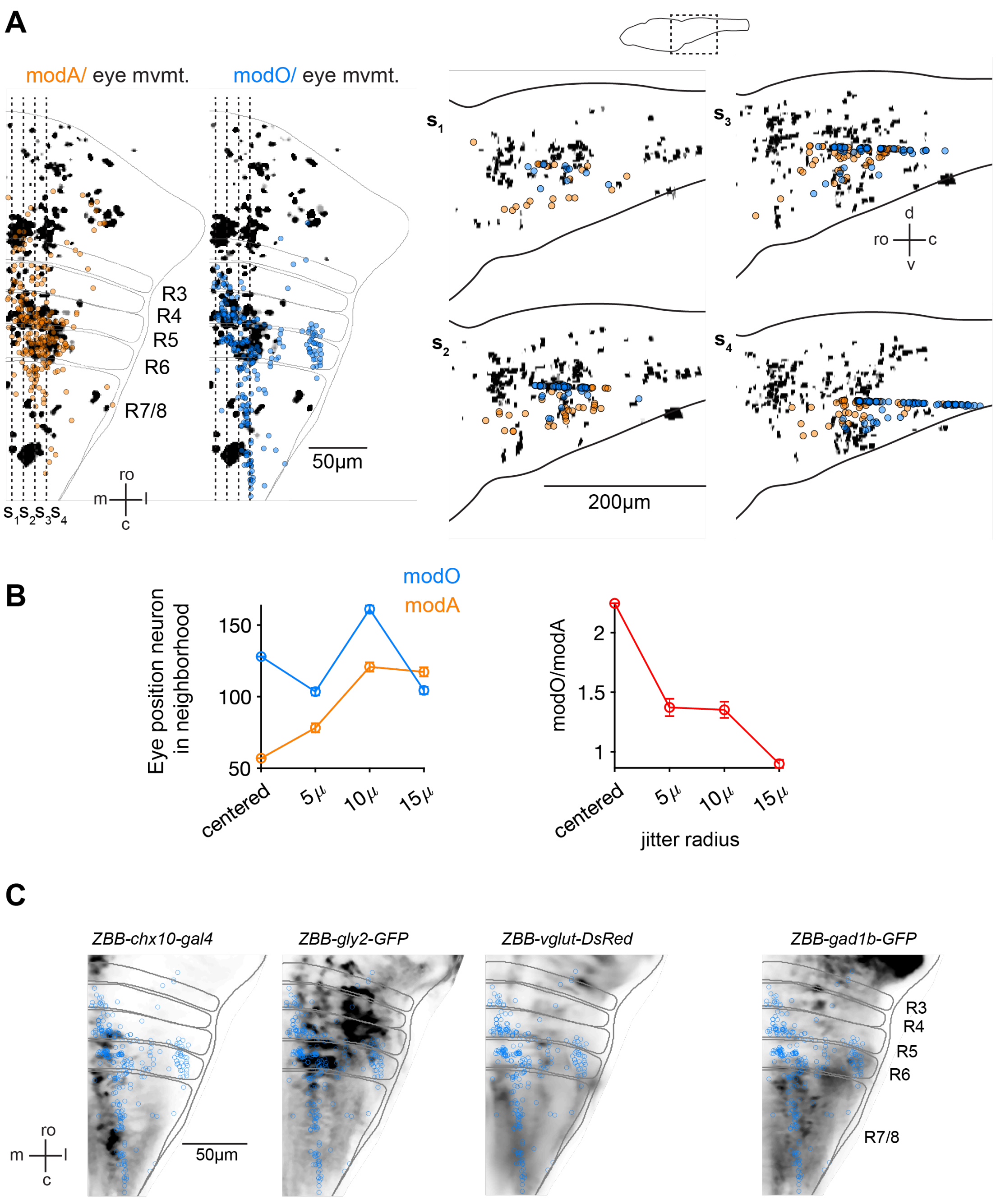


**Figure S9: Overlap of modules with functional maps.**
**A.** (Left) Overlap of neurons in modA (orange) and modO (blue) with a maximum intensity projection of neurons with eye movement signals (see Methods), functional map (background). Dotted lines indicate plane at which transverse views are visualized - (Right) Transverse views s1-s4 from left. Soma (foreground) and eye position signals (background) are over a span of 10$\mu$m along the medio-lateral axes centered on the dotted lines (ro - rostral; c - causal; m - medial; l - lateral; d - dorsal; v - ventral).
**B.** Quantification of the overlap of modO neurons from EM to neurons with eye movement signals functional imaging. (Top) Number of eye movement neurons as a function of jitter radius. Error bars are standard deviations of 10 iterations of jitter. (Bottom) ratio of eye position neurons as a function of soma jitter. Overlap was counted using a patch size of 6x6x8$\mu$m around each modO neuron.
**C.** Overlap of modO neurons registered to the Z-Brain atlas to transgenic lines that label neurotransmitter also in the same reference frame.

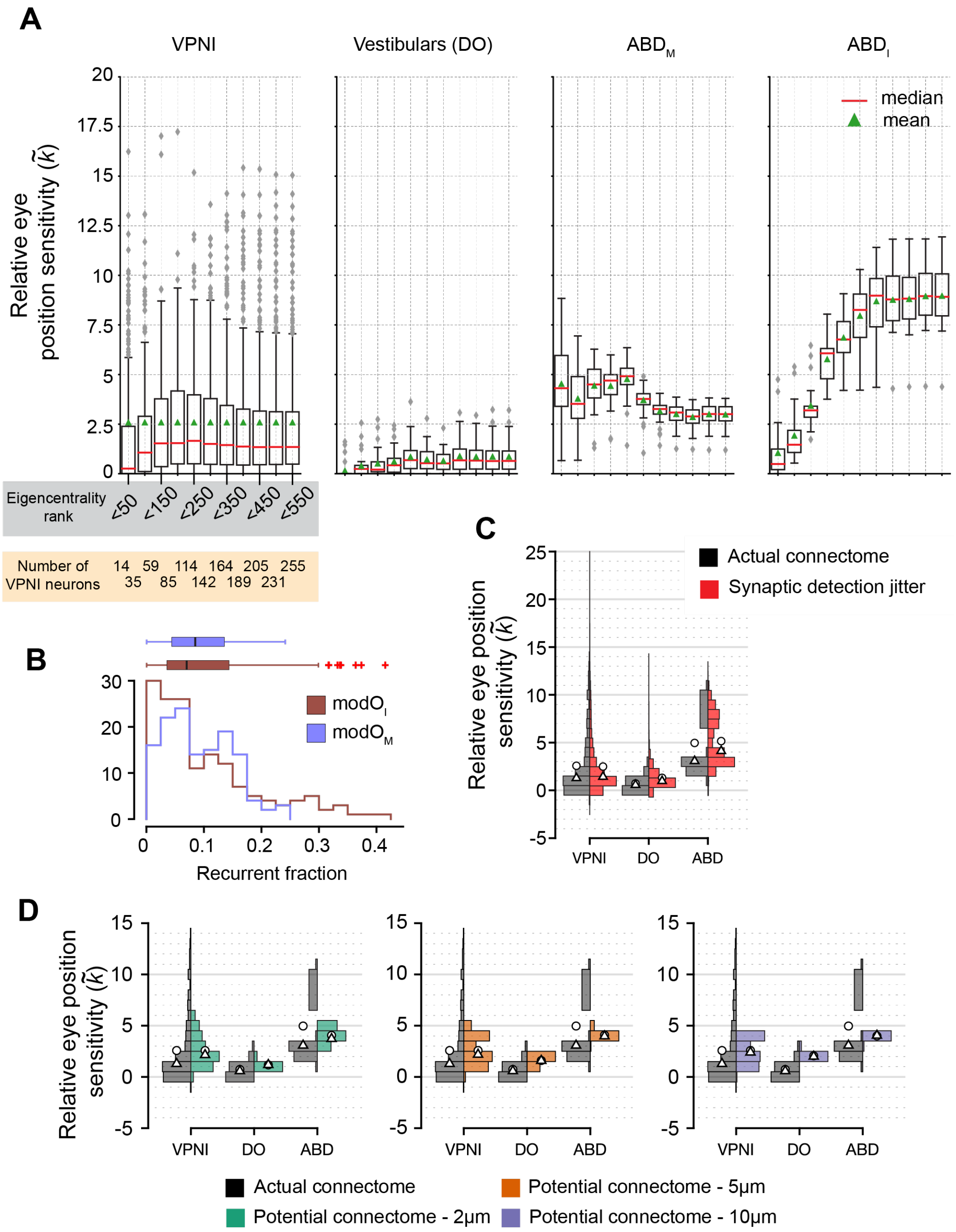


**Figure S10: Relative eye position sensitivity distributions for model variants.**

**A**. Box plots of relative eye position sensitivity $(\tilde{k})$ values for all four cell types, VPNIs, DOs , ABDm and ABDi neurons, when different numbers of neurons are used for the center. In these plots, the rightmost box represents the simulation shown in the main paper, when all center neurons are used. Boxes to the left of this indicate simulations run in which only neurons with a more stringent criterion for eigencentrality were included in the center, reducing the number of neurons in the population (number of VPNI model neurons included is shown in the highlighted box). The distributions of relative eye position sensitivity across the four neuronal populations remain consistent even when we shrink the candidate VPNI population by 50%.

**B.** Histogram of the fraction of all neurons that are recurrent within the identified modules. (left) Recurrent fraction within $modO_I$ and $modO_M$. Box plots, above, show medians along with $25^{th}$ and $75^{th}$ percentile. Red crosses are the outliers.

**C.** Distribution of the relative eye position sensitivities $(\tilde{k})$ when the model uses the actual connectivity matrix (black) compared to connectivity matrices generated by simulating small errors in automated synapse detection (red, see Methods). Circles represent the means and triangles represent the medians.

**D.** Distribution of the relative eye position sensitivities $(\tilde{k})$ when the model uses the actual connectivity matrix (black) compared to connectivity matrices generated considering potential synapses identified with increasingly poor resolution (green, brown, purple). Circles represent medians and triangles represent means. Simulations using connectomes with potential synapses quickly deviate from both those derived from the actual connectome and from the experimental distribution (Figure. 4E) as the potential synapse distance is increased.
