## supplemental information for "Predicting modular functions and neural coding of behavior from a synaptic wiring diagram"

### Supplementary Information

#### Gamification of the zebrafish dataset and crowdsourced reconstructions

In addition to in-house reconstructions by experts, zebrafish cells were made available to citizen scientist gamers through the existing retinal crowdsourcing game "Eyewire."

To test out this new dataset only the most experienced players on Eyewire were allowed to participate. A small group of 4 highly experienced players received an invite to test this new dataset. The players were given the title of "Mystic" which was a new status created to enable gameplay, and became the highest achievable rank in the game.

Subsequent Mystic players had to apply for the status to unlock access to the zebrafish dataset within Eyewire. There was a high threshold of achievement required for a player to gain Mystic status. Each player was required to reach the previous highest status within the game, as well as complete 200+ cubes a month and maintain 95% accuracy when resolving cell errors.

Once a player was approved by the lab, they were granted access to the zebrafish dataset, and given the option to have a live tutorial session with an Eyewire admin. There was also a pre-recorded tutorial video and written tutorial materials for players who could not attend the live session or who desired a review of the materials. Newly promoted players were also given a special badge, a new chat color, and access to a new chat room exclusive to Mystics. These rewards helped motivate players by showing their elevated status within the game as well as giving them a space to discuss issues specific to the zebrafish dataset.

Cells were parceled out in batches to players. When a cell went live on Eyewire, it could be claimed by any Mystic. Each cell was reconstructed by only one player at a time. Once this player had finished their reconstruction, a second player would check over the cell for errors. After the second check, a final check was done by an Eyewire Admin. To mitigate confusion when claiming cells, a special GUI was built into the game that allowed players to see the status of each cell currently online. A cell could be at one of five statuses - "Need Player A," "Player A," "Need Player B," "Player B," and "Need Admin." These statuses indicated whether a cell needed to be checked, or was in the process of being checked, and whether it needed a first, second, or Admin level check. At each stage the username of the player or Admin who had done a first, second, or final check was also visible. It was made mandatory that the first and second checks were done by two separate players.

Collaboration and feedback were important parts of the checking process. If a player was unsure about an area of the cell they were working on, they could leave a note with a screenshot and detail of the issue, or create an alert that would notify an Admin. If a "Player B" or an Admin noticed a mistake made earlier in the pipeline, they could inform the player of the issue via a "Review" document, or through an in-game notification (player tagging).

To differentiate the zebrafish cells from the regular dataset, each cell was labeled with a Mystic tag. This tag helped to identify the cells as separate from the e2198 retinal Eyewire dataset, and also populated them to a menu of active zebrafish cells within Eyewire.

Players were rewarded for their work in the zebrafish dataset with points. For every edit they made to a cell they received 300 points. Points earned while playing zebrafish were added to a player's overall points score for all gameplay done on Eyewire, and appeared in the Eyewire leaderboard.

<u>The following players in Eyewire were the admins who validated crowd sourced neuronal reconstructions:</u>

Hoodwinked, BenSilverman, Hightower, sarah.morejohn, SeldenK, sorek.m, zorek.m, twisterZ, hjones.jr, devonjones, amy, EinsteintheRapper, zkem, celiad, celiaz,sunreddy, peleaj43, sarahaw

<u>The following players helped reconstruct neurons on Eyewire, collectively called - Mystic players:</u>

r3, Atani, Nseraf, susi, eldendaf, Frosty, a5hm0r, hiigaran, kinryuu, Manni_Mammut, aesanta1, LotteryDiscountz, galarun, annkri, dragonturtle, LynneC, Cliodhna, jax123, KrzysztofKruk, Kfay, rinda, crazyman4865, JousterL, randompersonjci, Caffeine, Baraka, ggreminder, TR77, hewhoamareismyself, nagilooh, Oppen_heimer, Gruenewitwe, cognaso, twotwos, hawaiisunfun,danielag, lemongrab, zope, MysticM, kondor, frankenmsty, zfishman.
